# Drug resistant *Mycobacterium tuberculosis* strains have altered cell envelope hydrophobicity that influences infection outcomes in human macrophages

**DOI:** 10.1101/2024.04.10.588986

**Authors:** Alyssa Schami, M. Nurul Islam, Matthew Wall, Amberlee Hicks, Reagan Meredith, Barry Kreiswirth, Barun Mathema, John T. Belisle, Jordi B. Torrelles

## Abstract

*Mycobacterium tuberculosis* (*M.tb*), the causative agent of tuberculosis (TB), is considered one of the top infectious killers in the world. In recent decades, drug resistant (DR) strains of *M.tb* have emerged that make TB even more difficult to treat and pose a threat to public health. *M.tb* has a complex cell envelope that provides protection to the bacterium from chemotherapeutic agents. Although *M.tb* cell envelope lipids have been studied for decades, very little is known about how their levels change in relation to drug resistance. In this study, we examined changes in the cell envelope lipids [namely, phthiocerol dimycocerosates (PDIMs)], glycolipids [phosphatidyl-*myo*-inositol mannosides (PIMs)], and the PIM associated lipoglycans [lipomannan (LM); mannose-capped lipoarabinomannan (ManLAM)] of 11 *M.tb* strains that range from drug susceptible (DS) to multi-drug resistant (MDR) to pre-extensively drug resistant (pre-XDR). We show that there was an increase in the PDIMs:PIMs ratio as drug resistance increases, and provide evidence of PDIM species only present in the DR-*M.tb* strains studied. Overall, the LM and ManLAM cell envelope levels did not differ between DS- and DR-*M.tb* strains, but ManLAM surface exposure proportionally increased with drug resistance. Evaluation of host-pathogen interactions revealed that DR-*M.tb* strains have decreased association with human macrophages compared to DS strains. The pre-XDR *M.tb* strain with the largest PDIMs:PIMs ratio had decreased uptake, but increased intracellular growth rate at early time points post-infection when compared to the DS-*M.tb* strain H_37_R_v_. These findings suggest that PDIMs may play an important role in drug resistance and that this observed increase in hydrophobic cell envelope lipids on the DR-*M.tb* strains studied may influence *M.tb*-host interactions.

**AUTHOR SUMMARY:** Tuberculosis (TB) is a leading cause of death due to an infectious organism and is caused by the bacteria *Mycobacterium tuberculosis* (*M.tb*). Drug resistant (DR) forms of TB have emerged over the past few decades which make the disease even more difficult to diagnose and treat. Currently, there is very little known about how the bacteria changes as it becomes more drug resistant. Here, we used biochemical techniques to study differences in the *M.tb* cell envelope across drug resistance categories. We examined 11 *M.tb* strains that range from drug-susceptible (DS) to multi-drug resistant (MDR) to pre-extensively drug resistant (pre-XDR) and observed that levels of hydrophobic phthiocerol dimycocerosates were increased and levels of hydrophilic higher-order phosphatidyl-*myo*-inositol mannosides were decreased in DR-*M.tb* strains compared to DS strains. We also found that DR-*M.tb* strains had decreased association with human macrophages, and that the pre-XDR-*M.tb* strain with the highest ratio of hydrophobic to hydrophilic lipids had decreased uptake but increased intracellular growth in macrophages at early timepoints after infection. Our study provides exciting insights into changes in the cell envelope composition of DR-*M.tb* strains and how these changes may influence infection outcomes in human macrophages.

## INTRODUCTION

Tuberculosis (TB), caused by the organism *Mycobacterium tuberculosis* (*M.tb*), is considered one of the leading causes of death due to a single infectious organism with an estimated ∼1.3 million deaths annually.^1^ Recently, the prevalence of drug-resistant (DR) forms of TB such as multi-drug resistant (MDR), pre-extensively drug resistant (pre-XDR), and extensively drug resistant (XDR) has increased.^2^ This increase in DR-TB cases combined with the gap in knowledge related to DR-*M.tb* strains has made the TB disease even more difficult to diagnose and treat. To address DR-TB’s threat to public health, we must gain a greater understanding of drug resistance related changes in the *M.tb* cell envelope structure, and how these changes influence *M.tb*-host interactions.

The *M.tb* cell envelope is a dynamic and protective barrier and a target of various anti-TB drugs.^3^ Approximately 80% of the *M.tb* cell envelope is comprised of carbohydrates and lipids.^4^ The inner portion contains a plasma membrane and the covalently link cell wall ‘core’ that is sequentially comprised of peptidoglycan, arabinogalactan (AG) and the mycolic acid layers.^3–5^ The mycolic acid layer provides the foundation for the non-covalently intercalation of peripheral lipids, and together the mycolic acids peripheral lipids form the characteristic mycomembrane. Many of the peripheral lipids also act as virulence factors during infection and influence *M.tb*-host interactions.^3, 4^ However, there is limited information regarding how levels and the nature of these peripheral lipids change in relation to *M.tb* drug resistance.

Phthiocerol dimycocerosates (PDIMs) are aliphatic apolar peripheral lipids of the cell envelope of pathogenic mycobacterial species (with the exception of *M. gastri*).^6, 7^ Previous studies suggest that PDIMs suppress the early host immune responses by controlling TNF and IL-6 production and may also participate in arresting phagosomal maturation.^4, 8, 9^ PDIMs are also linked to cell envelope permeability and anti-TB drug susceptibility,^10–15^ although it is still unclear whether levels of PDIMs change as drug resistance increases. Conversely, phosphatidyl-*myo*-inositol mannosides (PIMs), especially the higher- order PIMs and their associated lipoglycans [namely, lipomannan (LM) and mannose-capped lipoarabinomannan (ManLAM)], are considered hydrophilic cell envelope molecules.^3, 4^ Previous studies examining PIMs, LM, and ManLAM found that they play multiple roles in *M.tb*-host interactions.^16–22^ In relation to drug resistance and virulence, data suggest greater variability in the size and branching of ManLAM in DR- and/or hypervirulent *M.tb* strains.^23–26^ There is also evidence that abolishment of LAM biosynthesis in *M. abscessus* is associated with increased susceptibility to multiple anti-TB drugs,^27^ and that treatment of *M.tb* with isoniazid (INH) results in the upregulation of genes related to ManLAM biosynthesis.^28^ Further, treatment of *M.tb* with ethambutol (EMB) results in truncated arabinan domains for both ManLAM and AG.^29, 30^ Initial studies investigated MDR- and XDR-*M.tb* strains using transmission electron and atomic force microscopy found increased cell envelope thickness of DR- compared to drug susceptible (DS)-*M.tb* strains.^31–35^ However, there are still limited data related to changes in specific components of the *M.tb* cell envelope as drug resistance profiles increase.

These findings led us to posit if levels of hydrophobic PDIMs and hydrophilic PIMs are changed in DR- *M.tb* strains compared to DS-*M.tb* strains. To test this, we biochemically examined the cell envelope composition of 11 different *M.tb* strains that ranged from DS to MDR to pre-XDR. Our findings indicate that the PDIMs:PIMs ratio is increased in all DR-*M.tb* strains studied. We provide evidence that the overall abundance of PDIMs increased gradually in our MDR- and pre-XDR strains, and in contrast the abundance of higher-order PIMs (but not LM or ManLAM) decreased in our DR-*M.tb* strains. We also provide evidence of PDIM species only present in the DR-*M.tb* strains studied. Further, we show that these *M.tb* strains studied had decreased association with primary human macrophages as their drug resistance profile increased, and that the pre-XDR strain with the highest PDIMs:PIMs ratio had decreased uptake but a faster intracellular growth rate at early timepoints after infection when compared to the DS strain H_37_R_v_. These observations suggest a greater hydrophobic composition of the DR-*M.tb* strains’ cell envelope compared to DS-*M.tb* strains, and that these changes in DR-*M.tb* cell envelope lipids may be associated with decreased infectivity but higher growth rates within macrophages *in vitro*.

## MATERIALS AND METHODS

### Ethics statement and human subjects

Human subject studies were carried out in strict accordance with the US Code of Federal and Local Regulations (Texas Biomedical Research Institute/UT Health San Antonio IRB number 20170315SHU). Blood from healthy adults (aged 18-45 years old) was collected from both sexes without discrimination of race and ethnicity after written consent as described.^36^

### M.tb strains and growth conditions

Three DS-*M.tb* strains [H_37_R_v_ (ATCC# 25618) Erdman (ATCC# 35801), and CDC1551 (NIH/NIAID BEI Resources# NR-13649) were used for all biochemical experiments and association studies. H_37_R_v_ and Erdman *M.tb* strains are common reference laboratory strains, and CDC1551 is a clinical isolate that is considered hypovirulent but highly transmissible.^37^ Three MDR- and 5 pre-XDR-*M.tb* strains (classified as MDR and pre-XDR based on recent World Health Organization definitions^38^) were provided by Drs. Kreiswirth and Mathema, from a collection of over 35,000 clinical isolates. For ease, the name of each DR clinical isolate has been changed to “MDR-x” or “pre-XDR-x” in all figures. **Table 1** lists each strain’s name (as used in our figures), original name (as provided by Drs. Kreiswirth and Mathema), genotypic resistance profile, and drug resistance category. All *M.tb* strains were grown for 21 days on solid 7H11 media supplemented with oleic acid, albumin, dextrose, catalase (OADC) enrichment. Single bacterial suspensions were prepared as described. ^39, 40^

**Table 1.**
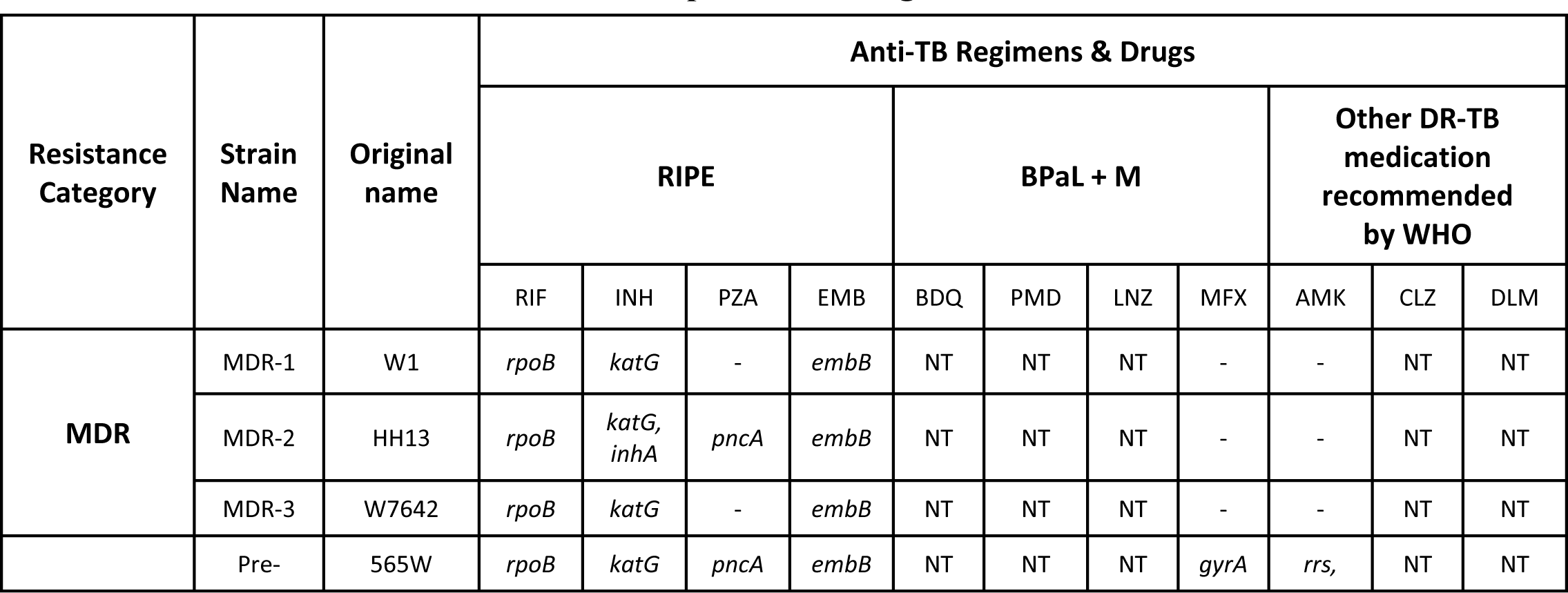
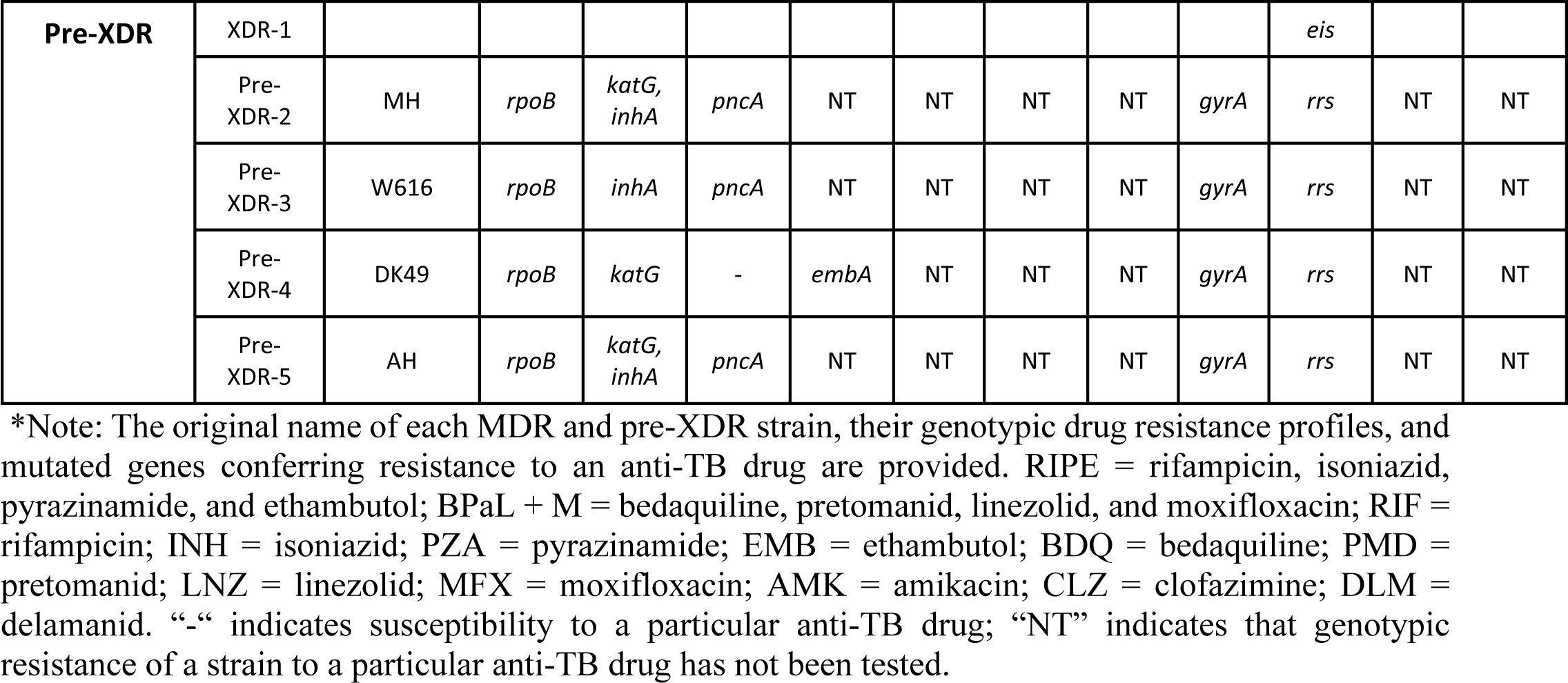
MDR- and Pre-XDR-*M.tb* strains studied and the mutated genes conferring their resistance to particular drugs *.

### Lipid extraction and analyses

Bacteria were harvested and then an exhaustive lipid extraction was performed. Briefly, *M.tb* was sequentially delipidated using chloroform:methanol 2:1 (*v/v*), chloroform:methanol 1:2 (*v/v*), and chloroform:methanol:water 10:10:3 (*v/v/v*), each for 24 h at 37°C. Lipid extracts from each organic solvent mixture were pooled together for total lipid analysis. For one-dimensional thin layer chromatography (1D-TLC) analysis of PDIMs, total crude lipids from each strain (normalized by biomass, equivalent to ∼100 µg of bacteria) were loaded onto 10 cm x 20 cm TLC silica gel 60 F_254_ aluminum plates and run in petroleum ether:acetone (96:4, *v/v*, run 3x). For two-dimensional TLC (2D-TLC) analysis of PIMs, total crude lipids from each strain (normalized by biomass; equivalent to ∼100 µg of bacteria) were loaded onto 10 cm x 10 cm TLC silica gel 60 F_254_ aluminum plates and run in the first dimension using chloroform:methanol:water (60:30:6, *v/v/v*) as a solvent system. The TLC plate was then rotated 90° to the left and run in the second dimension using chloroform:acetic acid:methanol:water (40:25:3:6, *v/v/v/v*) as a solvent system.^41^ TLC plates were dried and charred with 10% concentrated sulfuric acid in absolute ethanol and heated at 110°C for lipids visualization. NIH ImageJ program (http://rsb.info.nih.gov/ij/) was used to perform PDIMs and PIMs densitometry analyses by calculating mean spot intensities from four independent experiments. An independent experiment equates to new bacterial growth on 7H11 + OADC medium, new exhaustive lipid extractions, and subsequent TLC and densitometry analysis for each strain studied.

Liquid chromatography-mass spectrometry (LC-MS) analysis was also performed on the total lipid extract of each strain. Specifically, an aliquot (2 µg) of total lipid was applied to a XBridge C18 column (2.1 × 150 mm, 2.5 µm (Waters Corp.; Milford, MA) held at 45°C and that was attached to an Agilent 6546 liquid chromatography/quadrupole time-of-flight (LC/Q-TOF) mass spectrometer instrument (Agilent Technologies; Palo Alto, CA). The system was equilibrated with 100% solvent A [5 mM ammonium acetate in methanol]. Solvent A was maintained in an isocratic condition at 100% for 3.5 min, followed by a 30.0 min linear gradient to 98% solvent B [5 mM ammonium acetate in n-propanol-hexane (80:20; *v/v*)], and held at 98% solvent B for 3.0 min. Post-analysis time was set at 5 min to re-equilibrate the LC- MS system for the next sample analysis. The LC eluent was directly introduced into the Agilent 6546 Q- TOF mass spectrometer equipped with an electrospray ionization source. Q-TOF-MS operating parameters were: drying gas temperature at 310°C; sheath gas temperature at 270°C; drying gas flow at 8 L/min; sheath gas flow at 10 L/min; nebulizer pressure at 40 lb/in^2^; capillary voltage at 3500 V; and fragmentor voltage at 135 V. Positive (+) and negative (-) ion data were generated and mass spectra were acquired over a scan range of mass to charge (*m/z*) 250 to 3,000 at a scan rate of 1.25 spectra/sec. PDIM ion species were isolated in quadrupole for targeted MS/MS experiment at collision energy of 30 V. MS/MS data were acquired over a scan range of *m/z* 100 to 1,700 at a scan rate of 1.5 spectra/sec. All solvents and chemicals purchased were MS or HPLC grade. LC-MS instrument control and data acquisition were ascertained using the Agilent MassHunter WorkStation Data Acquisition software. PDIMs were analyzed in positive mode using Agilent MassHunter Qualitative Analysis 10 Software. PIMs were analyzed in negative mode using Skyline, an open-source Windows software system (https://skyline.ms/project/home/software/Skyline/begin.view), to extract the peak area values for each PIM based on its molecular formula and calculated the *m/z* ratio.

### Mannose-capped lipoarabinomannan (ManLAM) and lipomannan (LM) extraction and analysis

Delipidated bacterial pellets (normalized by biomass, equivalent to ∼100 μg of bacteria) from each *M.tb* strain underwent a quick extraction procedure to obtain ManLAM and LM as described.^42^ In brief, water and water-saturated phenol (1:1, *v/v*) were added to the exhaustively delipidated pellet and this mixture was incubated at 95°C for 2 h. Chloroform was added after 2 h followed by centrifugation at 10,000 x *g* at room temperature. The aqueous supernatant containing mainly ManLAM and LM was then dialyzed against water overnight using 3,500 molecular mass cutoff (MMCO) dialysis tubing. Extracts were analyzed by SDS-PAGE using 10-20% gradient Tris/Tricine gels followed by periodic acid-silver staining as described.^25, 42^ NIH ImageJ program (http://rsb.info.nih.gov/ij/) was used to perform ManLAM and LM densitometry analysis by calculating mean spot intensities from three independent experiments.

### Whole cell ELISA for ManLAM surface exposure

The surface exposure of ManLAM on each *M.tb* strain was assessed as described.^25^ Briefly, live *M.tb* single bacterial suspensions (1×10^6^) were added to each well (triplicate wells) of a 96-well tissue culture plate and dried at room temperature inside a biosafety cabinet. Wells were blocked with 1% bovine serum albumin (BSA) in phosphate-buffered saline (PBS) for 2 h. Wells were washed with PBS-T (1x PBS + 0.05% Tween-20) and incubated with either anti-LAM CS-35 (BEI Resources; Catalog # NR-13811) or CS-40 (BEI Resources; Catalog #NR-13812) mAbs in 1% BSA in PBS overnight at 4°C. Wells were washed with PBS-T and incubated with a secondary horseradish peroxidase-goat anti-mouse antibody (Abcam Ab 97023) for 2 h at room temperature. Wells were washed with PBS-T and developed using a TMB substrate kit (BD Biosciences; Catalog #555214). Reactions were stopped with 2M sulfuric acid and absorbance was measured at OD_450 nm_. Two independent experiments were performed, and an average value of absorbance from three triplicate wells for each *M.tb* strain was calculated per experiment.

### Human Monocyte-derived macrophage (hMDM) culture

Human peripheral blood mononuclear cells (PBMCs) were isolated from individual adult healthy donors using Ficoll-Paque cushion centrifugation, and cultured in Teflon wells with 20% autologous serum as described.^43^ On day 6, cells were harvested and hMDMs isolated by adherence to tissue culture plates for 2 h in RPMI 1640 medium + 10% autologous serum following the formation of confluent monolayers in 24-well plates containing sterile and treated coverslips (for association studies) or directly into the wells [for colony forming unit (CFU) studies]. hMDM monolayers were washed 3 times to remove lymphocytes and hMDM monolayers were incubated further overnight in RPMI 1640 + 10% autologous serum.

### Association and bacterial burden in infected hMDMs

On day 7, *M.tb* infections were performed where hMDM monolayers were infected with single bacterial suspensions of each *M.tb* strain at a multiplicity of infection (MOI) of 10:1 (for association studies) or 1:1 (CFU enumeration as previously described.^43^ Briefly, single cell suspensions of *M.tb* strains in RHH (RPMI 1640 + 10 mM HEPES + 0.1% human serum albumin) media (bacteria inoculum) were added to hMDM monolayers in coverslips at MOI 10:1. Plates were incubated on a shaker for 30 min and then incubated at rest for a total incubation time of 2 h at 37°C, 5% CO_2_. After infection, unbound bacteria were removed by washing several times with warm RPMI 1640, and fixed with 4% paraformaldehyde for 15 min at room temperature to assess for association by microscopy.

Similarly, hMDM monolayers in wells were infected with a MOI of 1:1 to determine intracellular growth at 2, 24, 72, and 120 h post-infection (hpi) using the CFU method as described.^39, 44^ In all experiments, synchronized infections were performed to ensure that all *M.tb* strains studied were in contact with the hMDM monolayer at the same time. This was achieved by gently centrifuging the hMDM monolayers at 300 x *g* at 4°C for 5 min during the process of initial infection and then, hMDM monolayers where further incubated for 2 h at 37°C, 5% CO_2_. After the 2 h infection time, for the CFU studies, monolayers were washed to remove unbounded bacteria, and further incubated with RMPI 1640 + 2% autologous, and CFUs assessed in each time point studied.

### Bacterial association measured by microscopy

Infected monolayers in coverslips were assessed for bacterial association by microscopy (bacteria per macrophage). Briefly, at 2 hpi, hMDM monolayers in coverslips were washed to remove unbound bacteria and fixed with 4% paraformaldehyde for 15 min at room temperature. To evaluate the *M.tb* association with hMDMs, coverslips were stained for bacteria using Auramine/Rhodamine (A/R) staining. hMDM nuclei were stained with 50 ng/ml 4’,6-diamidino-2- phenylindole (DAPI) (Invitrogen Cat. # D1306) for 7 min at room temperature. After multiple washes to remove the excess DAPI stain, coverslips containing stained *M.tb*-infected hMDM monolayers were mounted on slides using ProLong Gold Antifade Reagent (Invitrogen Cat. #P36934). Bacterial association per hMDM was evaluated with a Zeiss LSM 800 Confocal Microscope. For *M.tb* association, each A/R stained *M.tb* strain was quantified by examining the location of the bacteria in relation to the outer membrane of the hMDMs as was visualized using differential interference contrast (DIC) and fluorescence microscopy. Counting of bacteria for at least ≥ 300 consecutive hMDMs for each *M.tb* strain studied per human blood donor was performed as described.^40^ All strains tested, independent of their cell envelope composition and formation, stained similarly with A/R. All microscopy data were analyzed with Zeiss ZEN Software.

### Statistical analysis

GraphPad Prism 8 Software was used to prepare the graphs and determine the statistical significance between experimental groups. Student’s *t*-tests (unpaired) or One-way ANOVA with Dunnett’s post hoc correction for multiple comparisons were applied where applicable and as described in figure legends. In some cases, data sets that did not meet the assumption of normality were log transformed. Transformed data sets that still did not meet the assumption of normality were analyzed using Kruskal-Wallis nonparametric test and Dunn’s test for multiple comparisons. In this study, the “n” values for all biochemical experiments represent the number of biological replicas performed (*i.e.* the number of times a *M.tb* strain was grown and lipid or LAM/LM extraction was conducted). The “n” values for all cell biology experiments represent the number of human blood donors used as described in figure legends. Statistical differences between groups were reported as significant (*) when the *p*-value was less than or equal to 0.05.

## RESULTS

### DR-*M.tb* strains have increased PDIMs compared to DS-*M.tb* strains

Previous studies suggests that PDIMs may be a critical component to protect the impermeability of the *M.tb* cell envelope, as PDIMs deficiency is linked to increased susceptibility of mycobacteria to anti-TB drugs.^3, 11–13^ However, there are very few studies directed at the PDIMs levels in DR-*M.tb* strains.^15^ Here, we performed an exhaustive lipid extraction on three DS-, three MDR-, and five pre-XDR-*M.tb* strains and analyzed their total lipid extracts using TLC with validation by LC-MS. Our TLC analysis showed that PDIMs levels increase as the drug resistance profile increases, where pre-XDR *M.tb* strains had the largest increase in PDIMs levels when compared to DS-*M.tb* strains H_37_R_v_, Erdman, and CDC1551 (**Fig. 1A-B**; **Suppl. Fig. S1A**). Our LC-MS analysis detected 35 of the 52 PDIM species annotated by Sartain *et al.* (**Suppl. Table 1**).^45^ This analysis also confirmed an increase in PDIMs levels with an increase in drug resistance, where pre-XDR *M.tb* strains showed a significant increase in PDIMs levels *vs.* DS-*M.tb* strains (**Fig. 1C-D**; **Suppl. Fig. S1D**). These data are consistent with previous publications^8^ and indicate that while levels of PDIMs vary by strain (**Suppl. Fig. S1A** and **S1D**), there is an overall increase in PDIMs as the genotypic drug resistance profile increases.

**Figure 1.**
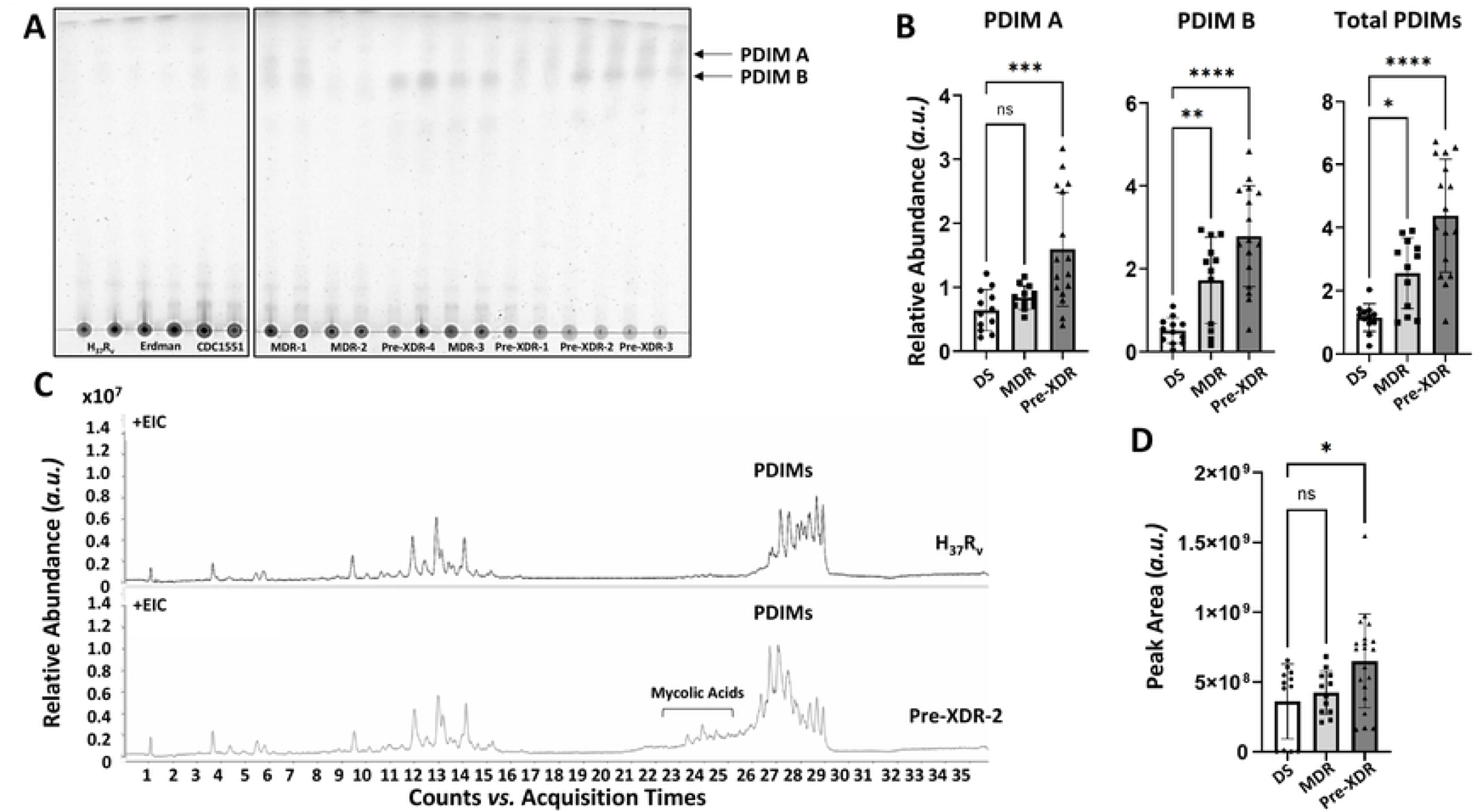
DR-*M.tb* strains have increased PDIMs compared to DS-*M.tb* strains. Lipids were obtained after an exhaustive organic solvent extraction and analyzed by 1D-TLC and LC-MS as described in our methods. **A)** Total lipids (normalized by biomass) were analyzed for PDIMs via 1D-TLC using Petroleum Ether:Acetone 96:4 (v/v, run 3x) as a solvent system and representative 1D-TLCs are shown (from n = 4 per strain). **B)** TLCs were quantified via densitometry analysis and compared across drug resistance categories for PDIM A, PDIM B, and total PDIMs. **C)** Total lipids (normalized by biomass) were analyzed for PDIMs via LC-MS analysis in positive mode. Representative extracted ion chromatograms (EIC) of H_37_R_v_ and Pre-XDR-2 are shown, with PDIMs eluting between 26 to 29 min. **D)** Peak area values for PDIMs [mass to charge (*m/z*) values between 1250-1540] were calculated using Agilent MassHunter Qualitative Analysis software and compared across drug resistance categories. Graphs are shown as M ± SD, where each data point represents an individual lipid extraction. N = 12-20 extracts per drug resistance category, where the “n” value represents the number of biological replicas performed (*i.e.* the number of times a *M.tb* strain was grown and lipid extraction was conducted). Graphs were analyzed using One-way ANOVA with post-hoc Dunnett’s test. In some cases, data sets that did not meet the assumption of normality were log transformed. Transformed data sets that still did not meet the assumption of normality were analyzed using Kruskal-Wallis nonparametric test and Dunn’s test for multiple comparisons. *p<0.05; **p<0.01; ***p<0.001.

### DR-*M.tb* strains have multiple PDIMs isomers that are not present in DS-*M.tb*

During our LC-MS validation of the lipid extracts, we uncovered that several of the DR-*M.tb* strains studied produce two isomers of three PDIM A and three PDIM B alkylforms (molecules of the PDIM family that differ in length and saturation of fatty acyl or polyketide backbones^46^) that were not present in the DS-*M.tb* strains (**Fig. 2**; **Tables 2** and **3**). Each PDIM isomer pair had the same molecular formula (and thus, the same *m/z* value), but different retention times. As shown in **Figure 2A** for the PDIM A species with an [M+NH_4_]^+^ molecular ion of 1399.446 *m/z* and a molecular formula of C_93_H_184_O_5_, the isomer present only the in DR-*M.tb* strains eluted with an earlier retention time compared to the isomer present in both the DS- and DR-*M.tb* strains. PDIM molecular ions were characterized by the formation of [M+Na]^+^ and [M+NH_4_]^+^ adducts, and the MS spectra underlying PDIM A (C_93_H_184_O_5_) isomers at 28.096 min and 28.119 min in the DS strain H_37_R_v_ and Pre-XDR-2 strain, respectively, and the peak at 27.053 min for the Pre-XDR-2 strain were identical (**Fig. 2B-D**). Similar LC-MS findings were obtained for each isomer pair (data not shown). Of note, these isomers were not detected for all PDIM alkylforms and were more abundantly detected in PDIM A when compared to PDIM B, except for MDR-2 and Pre- XDR-5 *M.tb* strains in which isomers only occurred for PDIM B (**Fig. 2E-F**; **Table 2** and **3**). Additionally, some DR-*M.tb* strains produced more of these isomers than other strains, and not every DR-*M.tb* strain produced each novel isomer (**Table 2** and **3**). Indeed, the MDR-2 *M.tb* strain produced low amounts of isomers related to the PDIM B alkylform C_95_H_186_O_5_, but no other isomers were detected for this strain (**Table 3**). Thus, here we included a table of the three most abundant alkylforms containing isomers for each *M.tb* strain studied for PDIM A (**Table 2**) and PDIM B (**Table 3**), and denoted which strain(s) did not contain specific isomers. Overall, these data suggest that DR-*M.tb* may have multiple PDIM isomers that are not present in DS-*M.tb* strains, which could contribute to increasing their hydrophobicity; however, more testing is necessary to determine the structure of these isomers and the mechanism behind their biosynthesis.

**Figure 2.**
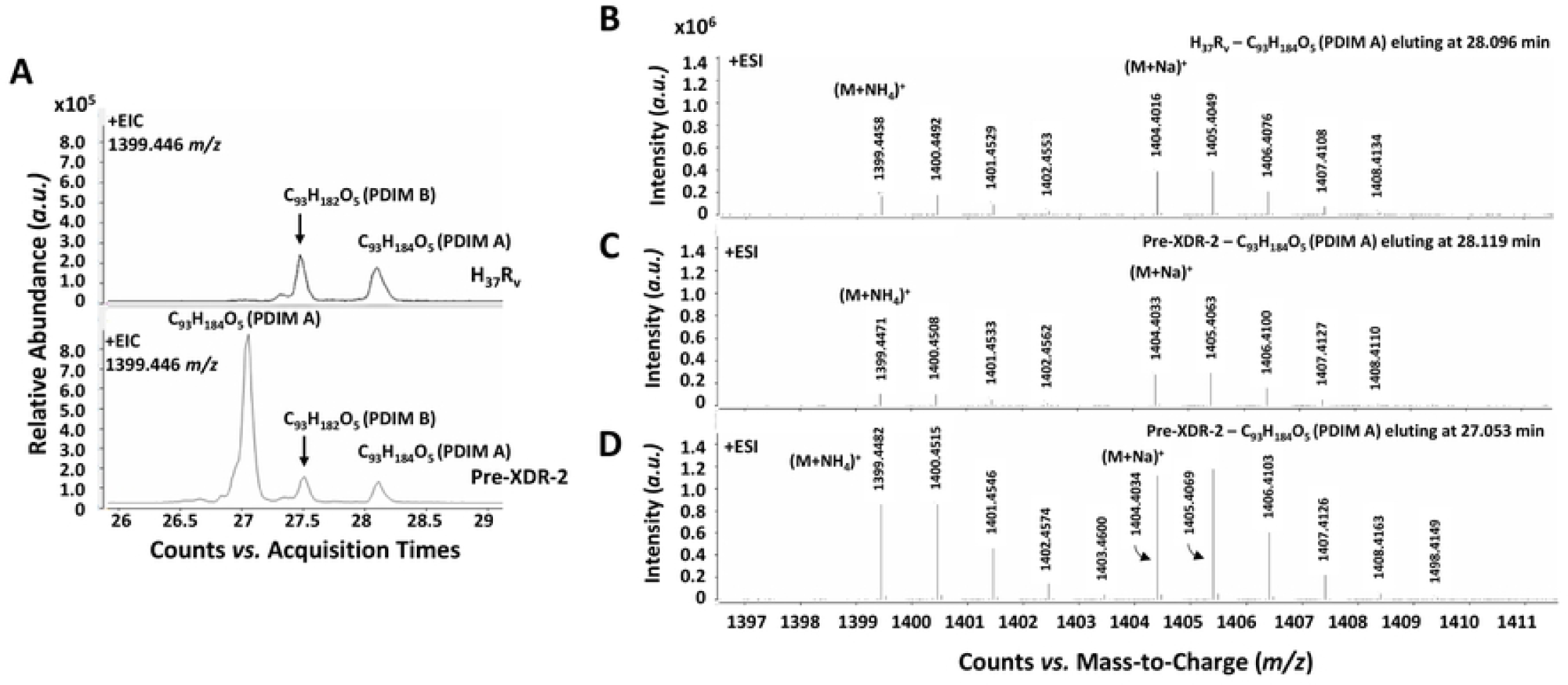
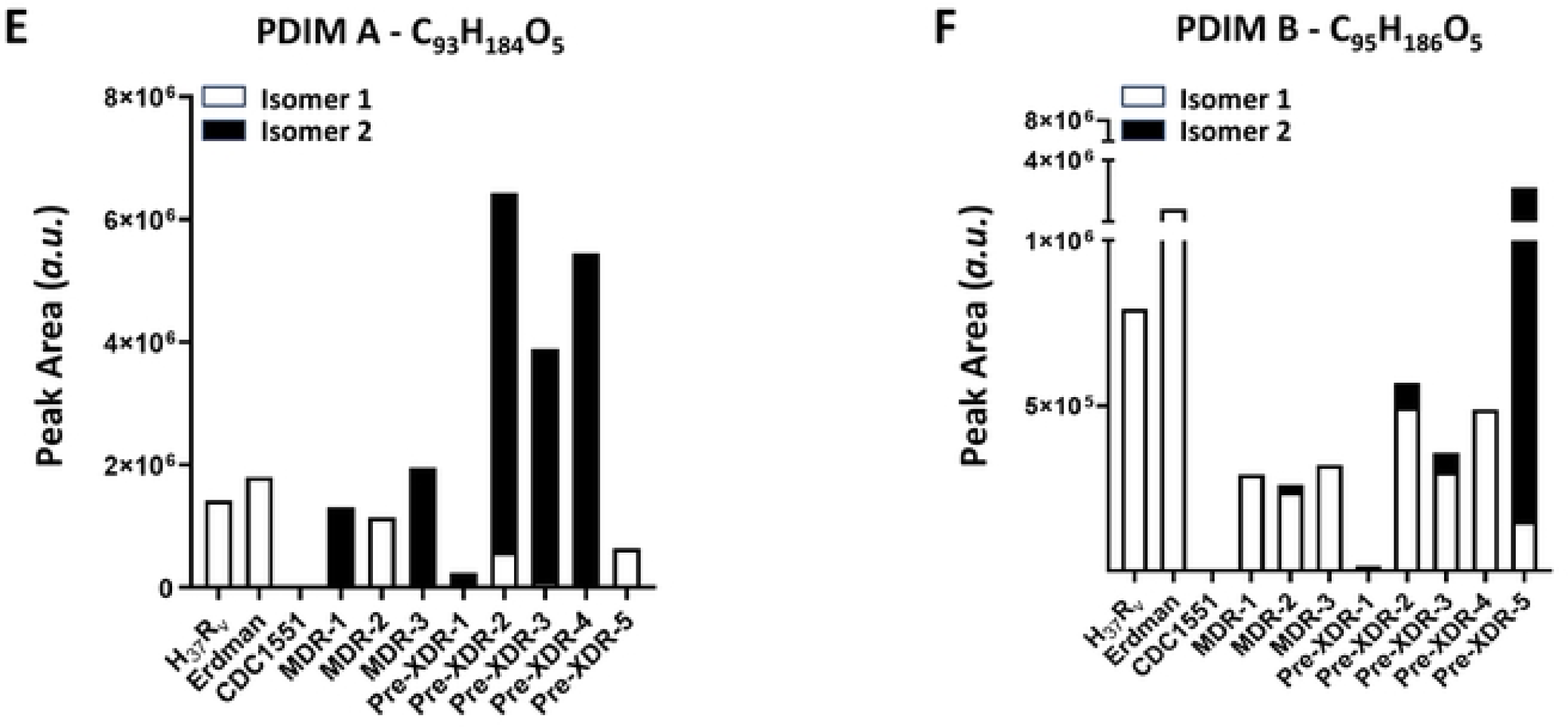
DR-*M.tb* strains have PDIM isomers that are not present in DS-*M.tb*. Lipids were obtained after an exhaustive organic solvent extraction and analyzed by LC-MS as described in our methods. Total lipids (normalized by biomass) were analyzed for PDIMs via LC-MS analysis in positive mode. **A)** Representative extracted ion chromatograms (EIC) of H_37_R_v_ and Pre-XDR-2 *M.tb* strains are shown for the PDIM A alkylform C_93_H_184_O_5_ (*m/z* of 1399.446). **B-D)** Mass spectra for the EIC peak of PDIM A C_93_H_184_O_5_ (*m/z* of 1399.446) at **B)** 28.096 min for *M.tb* H_37_R_v_, **C)** 28.119 min for *M.tb* Pre-XDR-2, and **D)** 27.053 min for *M.tb* Pre-XDR-2 are identical. It was noted that the EIC for 1399.446 *m/z* also resulted in a peak at 27.470 and 27.506 min in both the H_37_R_v_ and Pre-XDR-2 *M.tb* strains, respectively. However, the MS spectra at this retention time reveled that this was a ^13^C_2_ isotopic ion of the PDIM B species with a [M+NH_4_]^+^ of 1397.431 *m/z* (**See Suppl. Fig. S2**). **E-F)** Quantification of **E)** PDIM A C_93_H_184_O_5_ isomers and **F)** PDIM B C_95_H_186_O_5_ isomers for all *M.tb* strains are shown. Graphs are shown as the mean value of 4 biological replicates per strain for 1 or 2 isomers for each PDIM alkylform.

**Table 2.**
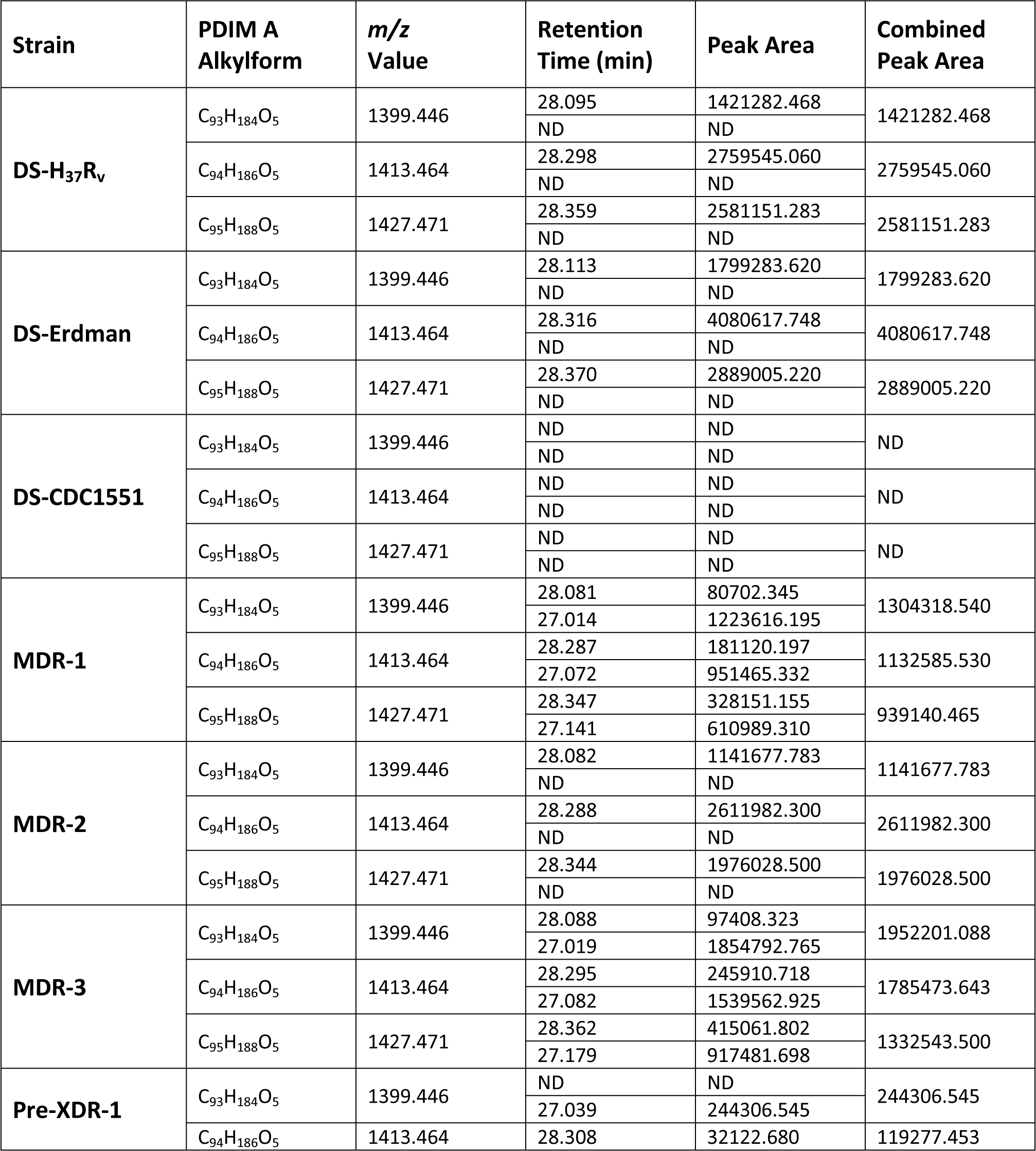
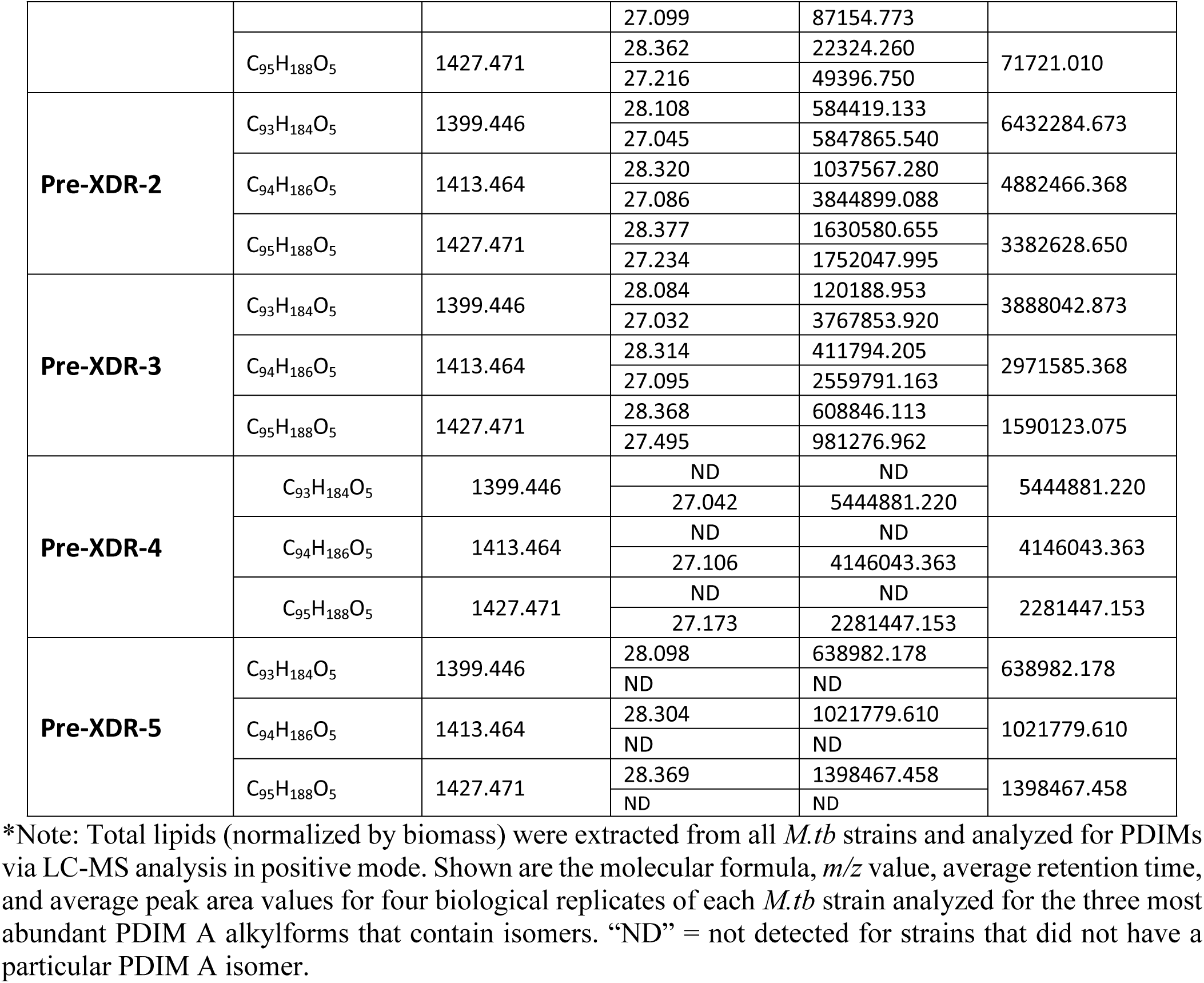
Abundance of PDIM A isomers in DR-*M.tb* strains*.

**Table 3.**
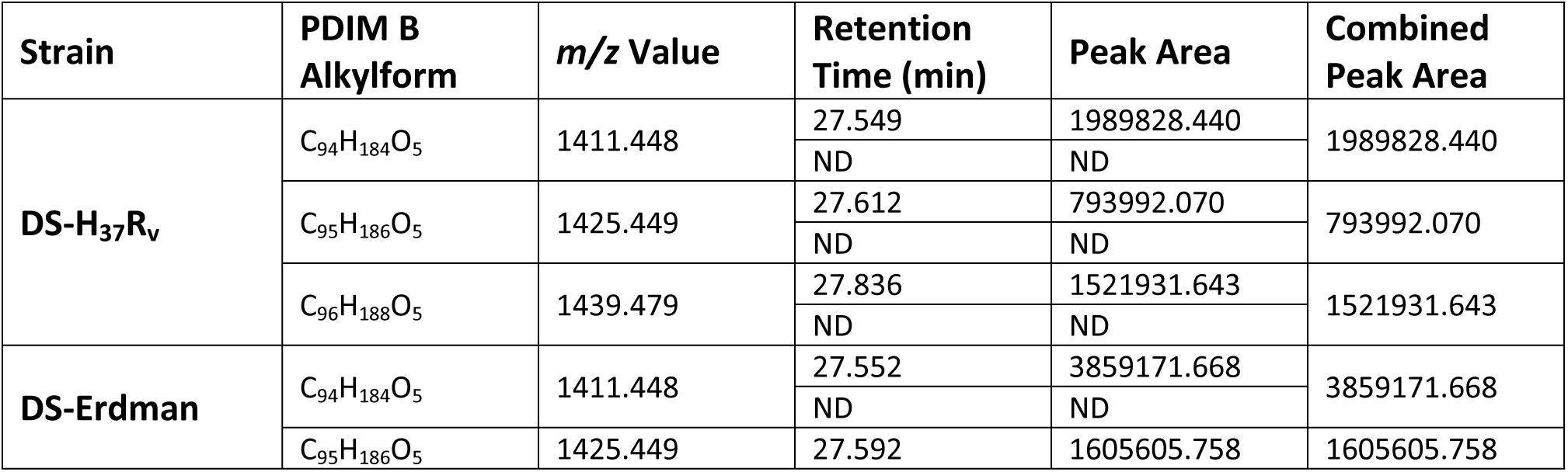
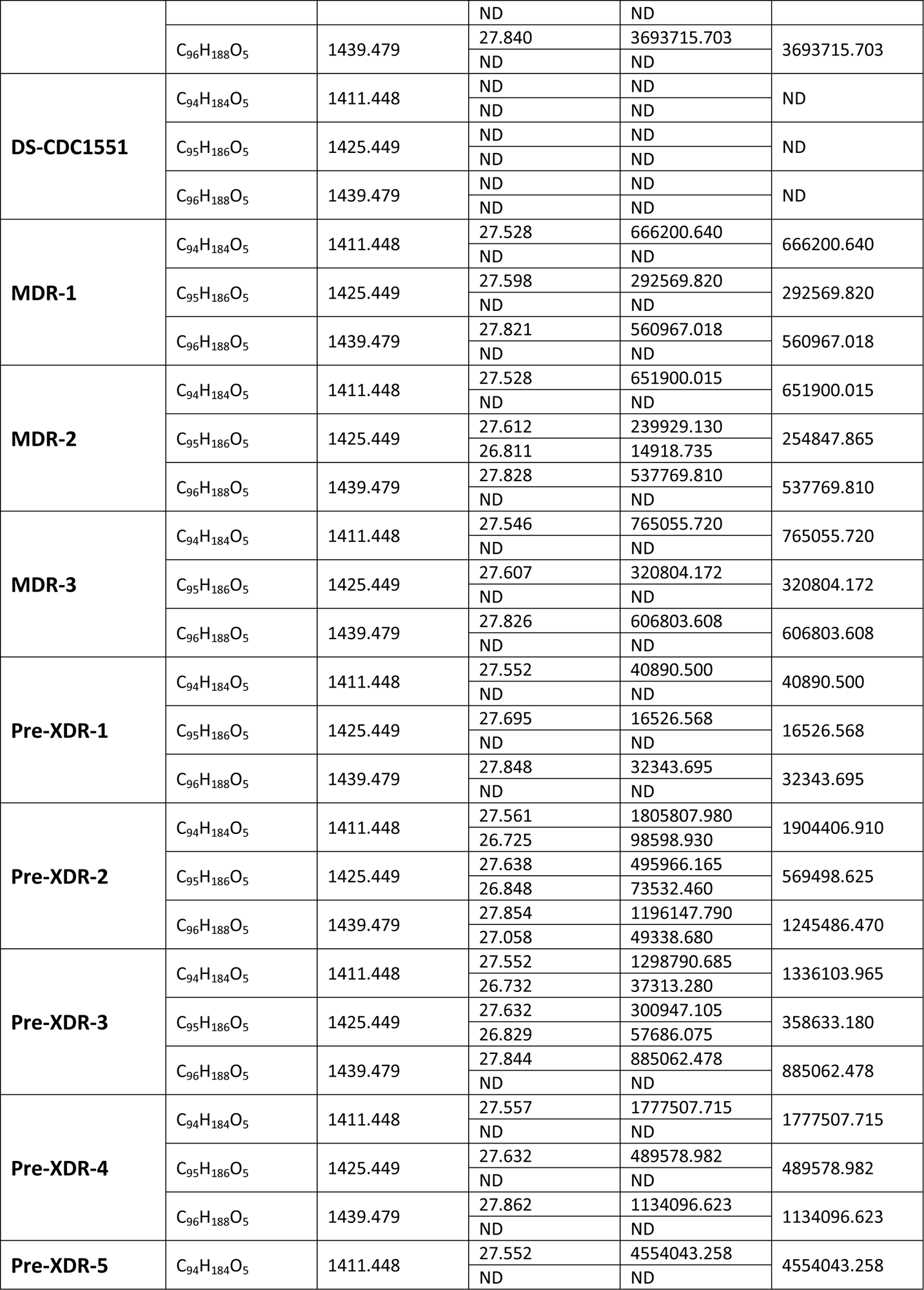
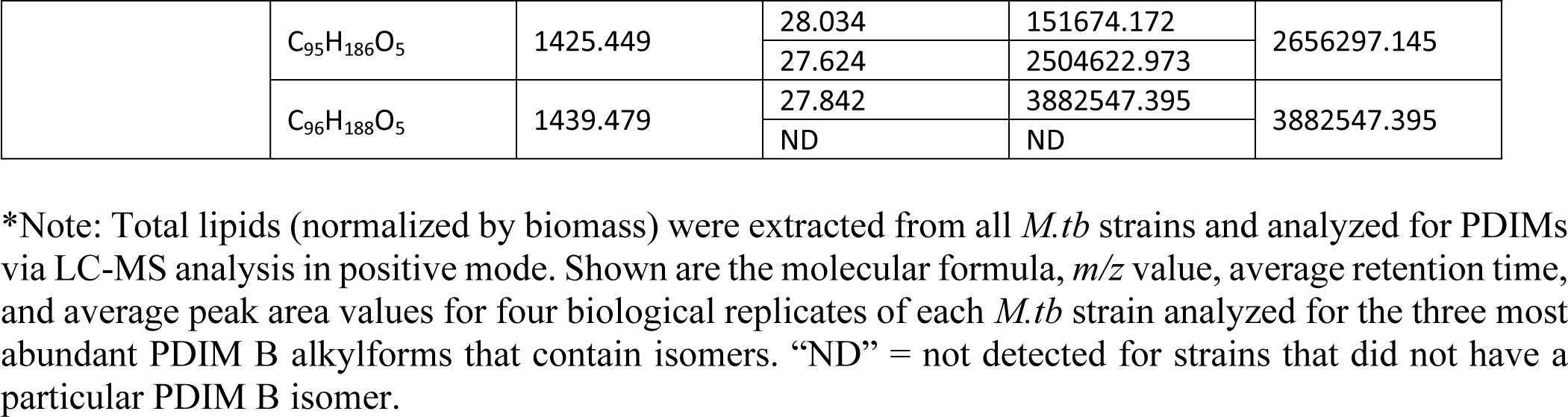
Abundance of PDIM B isomers in DR-*M.tb* strains*.

Recently, Flentie *et al.* mapped retention times and established MS/MS fragmentation patterns for the PDIM A and PDIM B species of *M.tb*.^47^ Using this as a guide, targeted MS/MS was carried out on [M+NH_4_]^+^ adduct of the isomers to identify characteristic fragments ions and define the structures. As shown in **Fig. 3A and 3B**, MS/MS spectrum of the *m/z* 1399.446 molecular ion of the PDIM A isomer that eluted at ∼ 28.1 min in the DS strain H_37_R_v_ and Pre-XDR-2 strain produced nearly identical patterns with major fragments ions at *m/z* 943.98 and 929.96 corresponding to the neutral loss of C29:0 and C30:0 mycocerosic acids plus ammonium from a phthiocerol back bone. Minor fragment ions at *m/z* 972.01 and 901.93 also indicated a minor species substituted with C27:0 and C32:0. In contrast the early eluting isomer (27.053 min) of the Pre-XDR-2 strain, yielded a MS/MS spectrum dominated by *m/z* 943.98 fragment ion, indicating a PDIM A species with two C29:0 mycocerosic acids (**Fig. 3C**). Minor fragment ions of *m/z* 986.02 and 901.83 corresponded to a minor PDIM A species with a C26:0 and C32:0 mycocerosic acid composition. The retention time difference of the isomers, however, cannot be attributed to slight differences in mycocerosic acid chain lengths. Flentie *et al.* showed a nearly 1 min difference in the retention time of PDIM A species with a phthiocerol (C3 methoxy) versus PDIM A species with a phthiodiolone (C3 keto) backbone, the latter yielding an earlier retention time.^47^ It is not possible to have isomeric species of PDIMs with a phthiocerol and a phthiodiolone backbone; however, the less commonly observed phthiotriol (C3 hydroxy) based PDIMs will result in an isomeric structure with the phthiocerol based PDIMs and would be expected to have an earlier retention time by reversed phase chromatography. Thus, we have concluded the early eluting PDIM A isomer associated with the DR strains are a result of a phthiotriol backbone instead of a phthiocerol.

**Figure 3.**
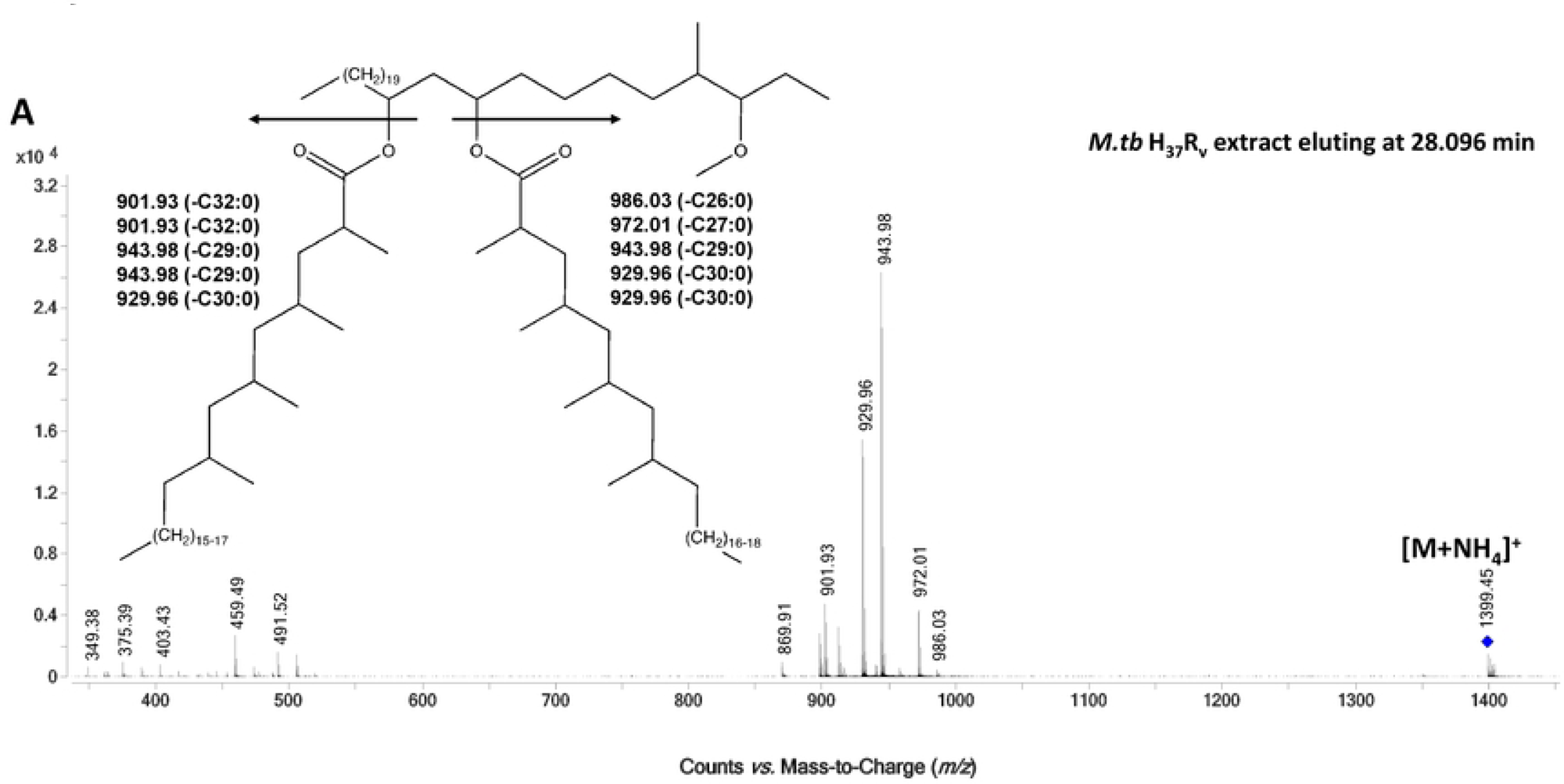
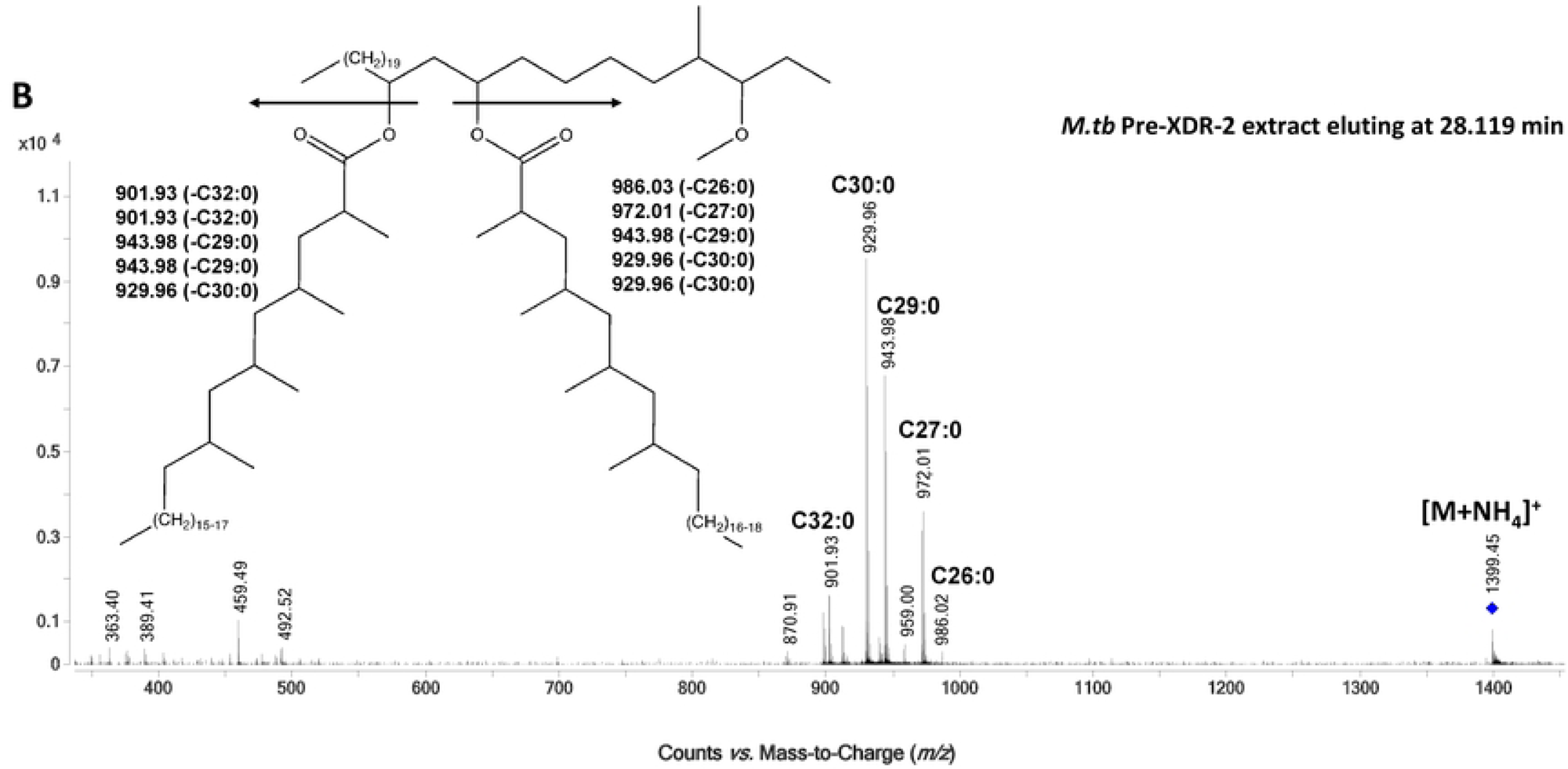
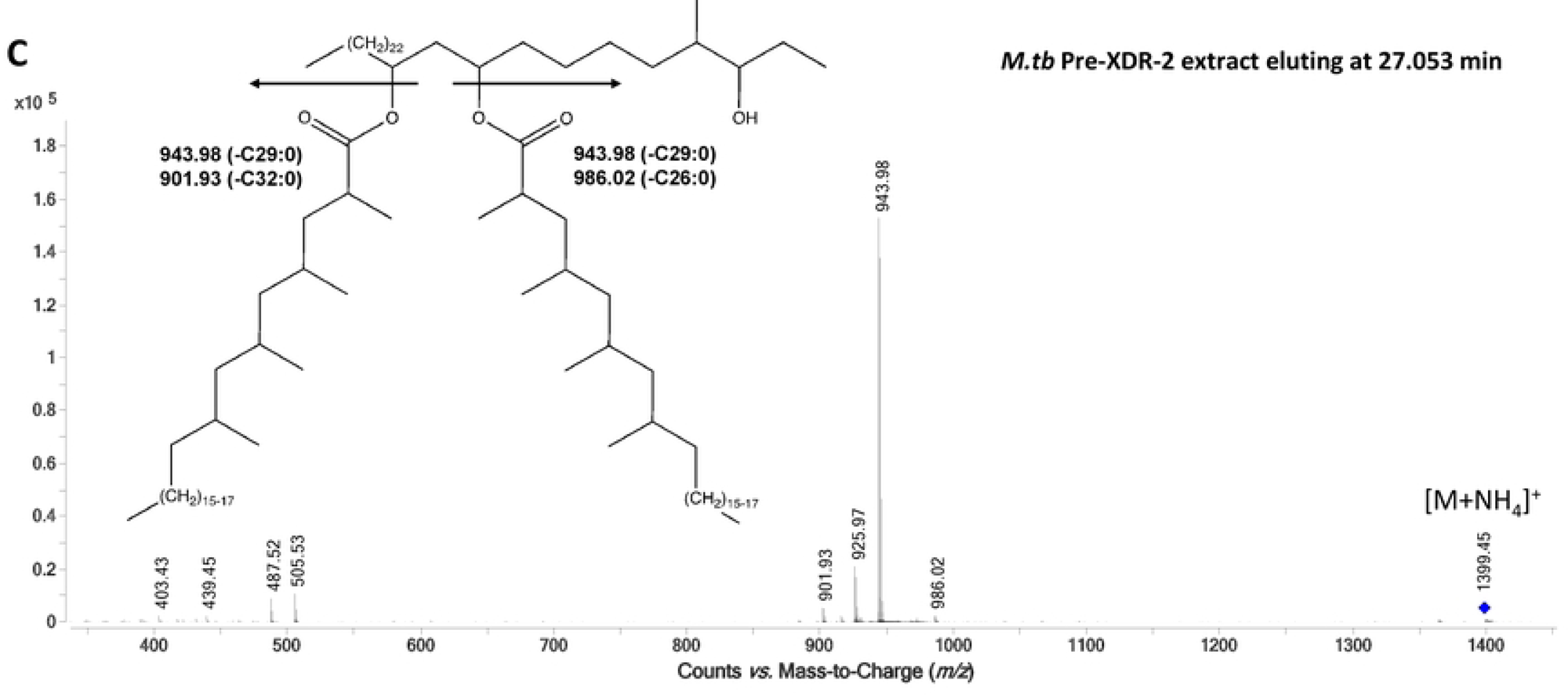
LC-MS/MS spectra of PDIM isomers in DS- and DR-*M.tb* strains. Lipids were obtained after an exhaustive organic solvent extraction and analyzed by LC-MS/MS as described in our methods. Total lipids (normalized by biomass) were analyzed for PDIMs via LC-MS analysis in positive mode, and PDIM isomers were analyzed using Agilent MassHunter Qualitative Analysis 10 Software. **A-C)** Representative MS/MS spectra of **A)** *M.tb* H_37_R_v_’s C_93_H_184_O_5_ (eluting at 28.096 min) and **B)** *M.tb* Pre- XDR-2 C_93_H_184_O_5_ isomer 1 (eluting at 28.119 min) revealed the same fragmentation pattern. **C)** Representative MS/MS spectra of Pre-XDR-2’s C_93_H_184_O_5_ isomer 2 (eluting at 27.053 min) showed a similar but not identical fragmentation pattern. Insets depict the corresponding PDIM structures and the fragment ions expected based on neutral loss of the mycocerosic acids with NH_3_.

### DR-*M.tb* strains have decreased PIMs compared to DS-*M.tb*

Our next question was whether levels of PIMs, hydrophilic mannose-containing glycolipids on the *M.tb* cell envelope, also change in DR-*M.tb* strains. Our analysis of PIMs using 2D-TLC and densitometry analysis indicated that total PIMs, lower-order, and higher-order PIMs levels were all decreased in both MDR and pre-XDR strains compared to DS-*M.tb* (**Fig. 4A-B**; **Suppl. Figs. S1B** and **S3A-B**). We further confirmed these 2D-TLC results by LC-MS analysis (**Fig. 4D-F**). While total PIMs still showed a slight decrease using LC-MS, we did not see a significant decrease in total PIMs in either MDR or pre-XDR resistance categories (**Fig. 4F**; **Suppl. Fig. 1E**). However, after separating total PIMs into lower- and higher-order PIMs, we were able to confirm that higher-order PIMs were significantly decreased in both MDR and pre-XDR *M.tb* strains when compared to DS *M.tb* strains (**Fig. 4D-F; Suppl. Fig. S3C-D**).

**Figure 4.**
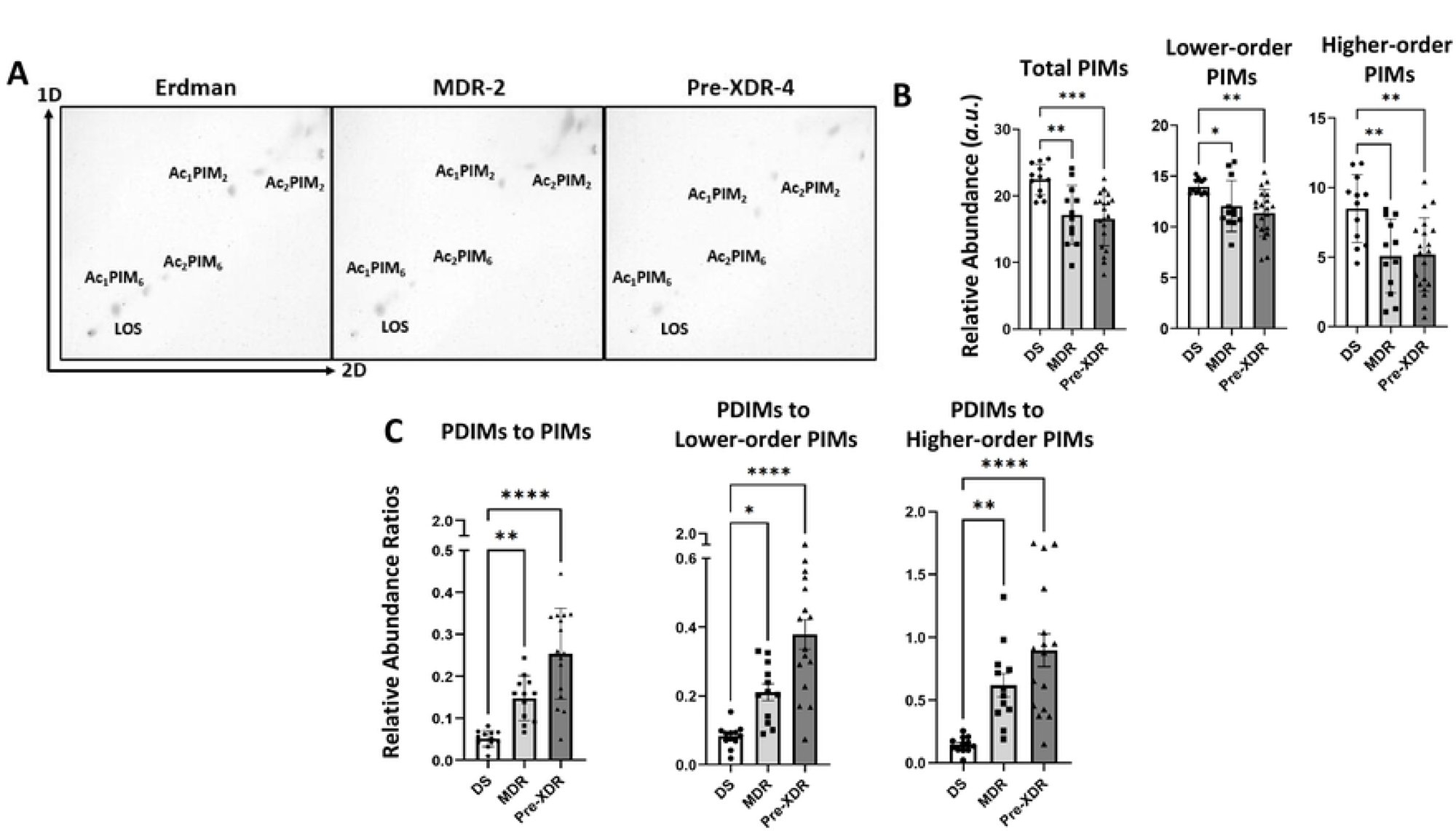
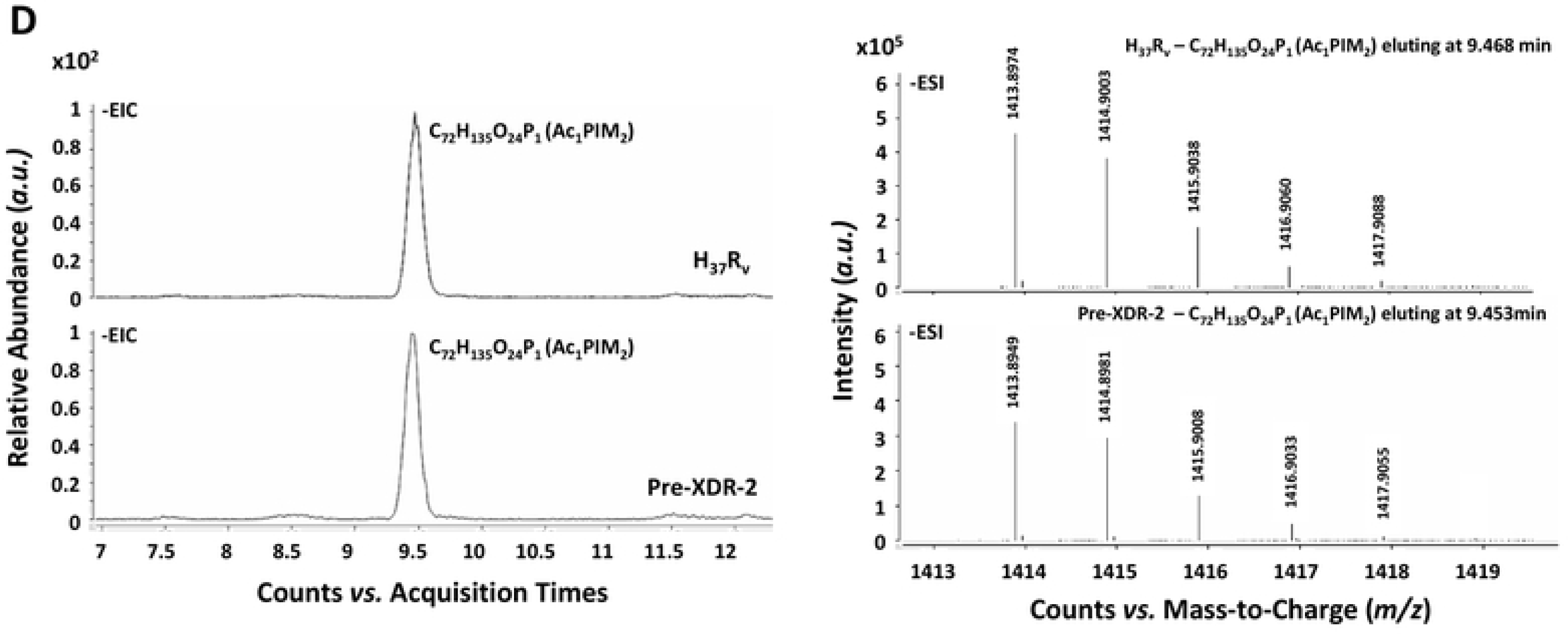
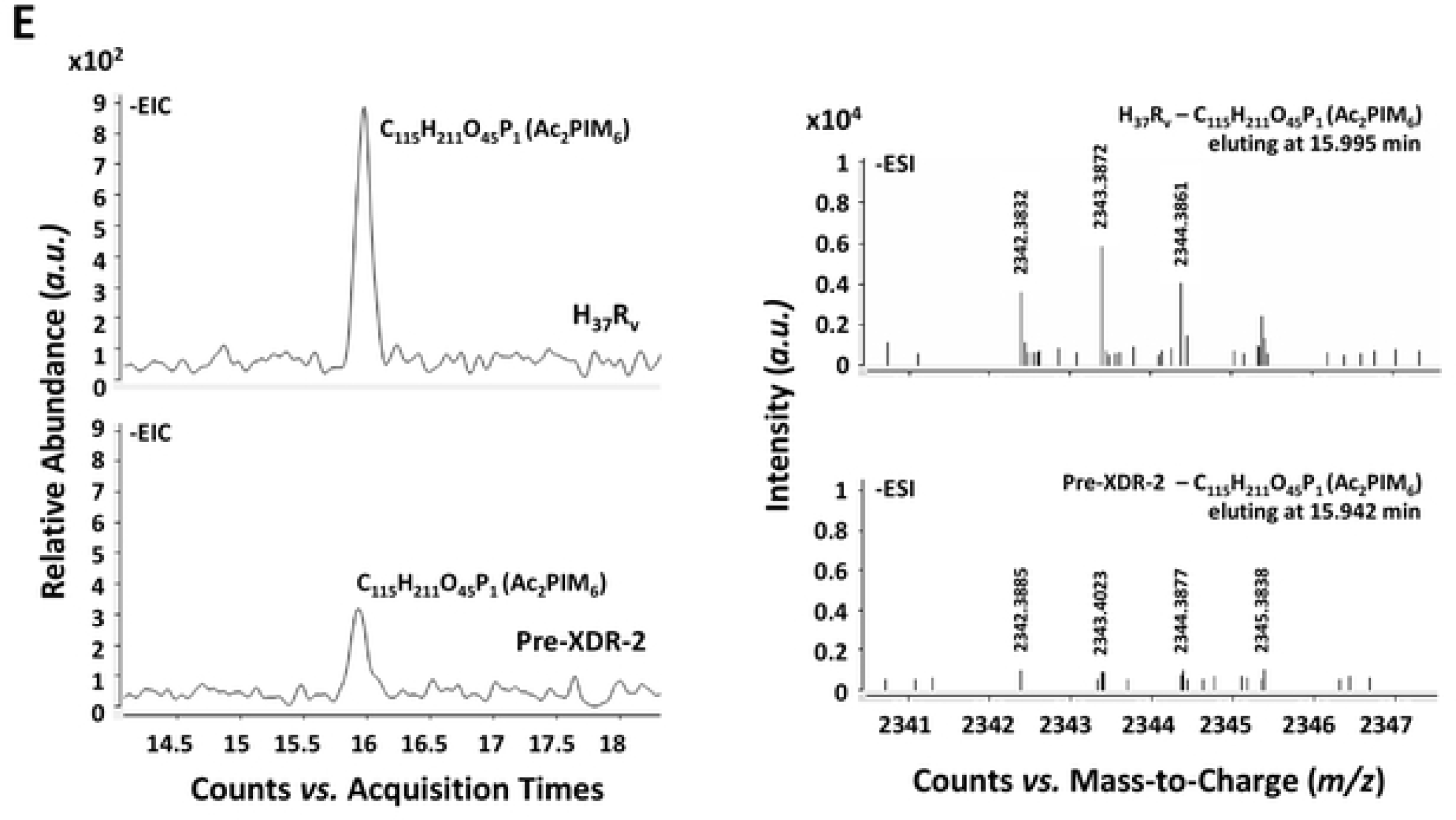
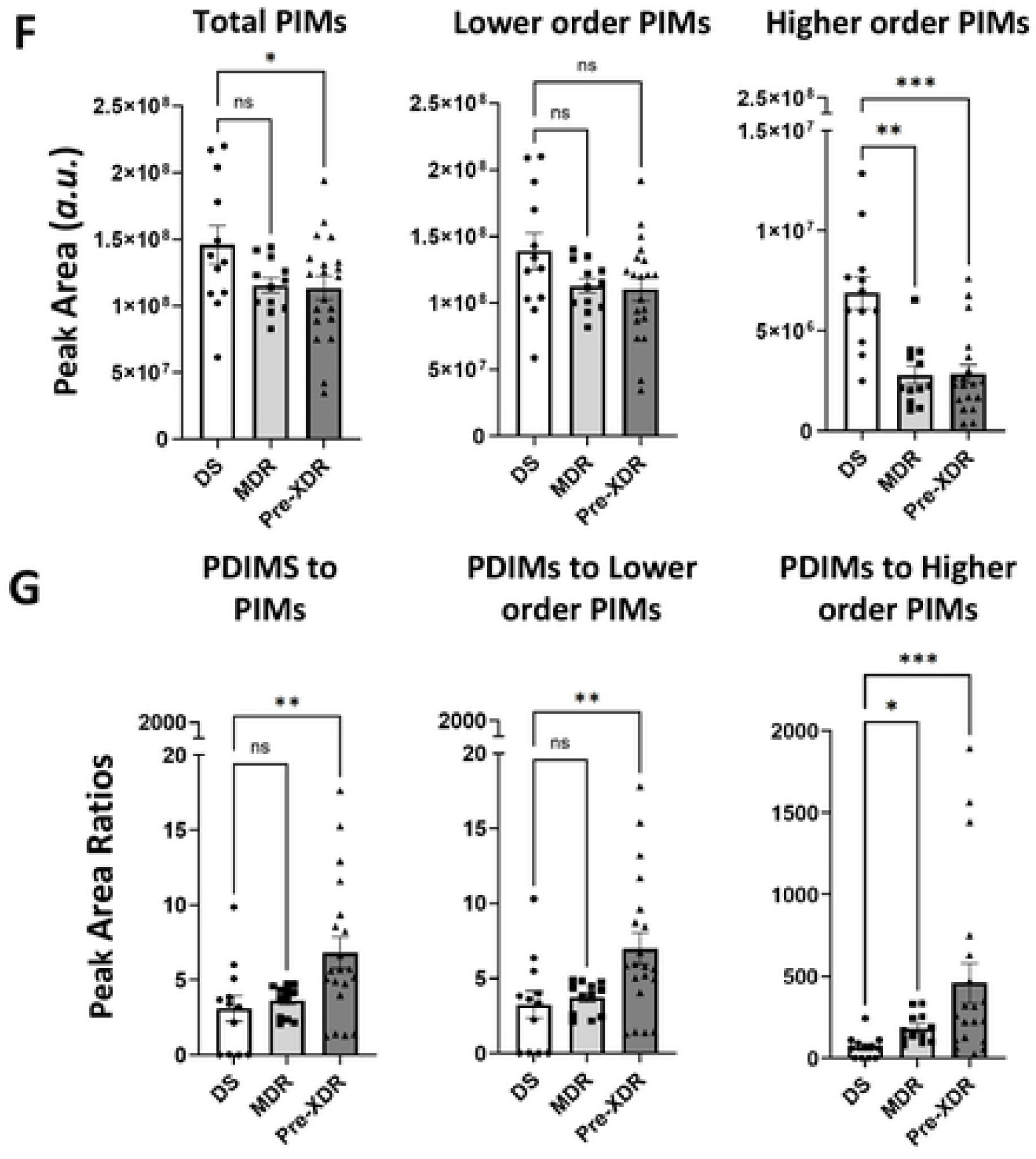
DR-*M.tb* strains have decreased higher-order PIMs compared to DS-*M.tb*. Lipids were obtained after an exhaustive organic solvent extraction and analyzed by 2D-TLC and LC-MS as described in our methods. **A)** Total lipids (normalized by biomass) were analyzed for PIMs via 2D-TLC using chloroform:methanol:water (60:30:6,v/v/v) in the first dimension (1D). TLC plates were then rotated 90° to the left and run using chloroform:acetic acid:methanol:water (40:25:3:6, v/v/v/v) in the second dimension (2D). Representative 2D-TLCs of DS, MDR, and pre-XDR *M.tb* strains are shown. **B)** 2D- TLCs were quantified via densitometry analysis and compared across drug resistance categories for total PIMs, lower-order PIMs, and higher-order PIMs. **C)** The relative abundance ratios of PDIMs to total PIMs, lower-order PIMs, and higher-order PIMs were calculated for each *M.tb* strain and compared across drug resistance categories. **D-F)** Total lipids (normalized by biomass) were analyzed for PIMs via LC-MS analysis in negative mode. **D)** Representative extracted ion chromatograms (EIC) of H_37_R_v_ and Pre-XDR- 2 *M.tb* strains for the lower-order PIM Ac_1_PIM_2_ and their respective mass spectra are shown. **E)** Representative extracted ion chromatograms (EIC) of H_37_R_v_ and Pre-XDR-2 for the higher-order PIM Ac_2_PIM_6_ and their respective mass spectra are shown. **F)** Peak area values for PIMs were calculated using Skyline software (using known retention times and *m/z* values of each PIM) in negative mode and compared across drug resistance categories for total PIMs, lower-order PIMs, and higher-order PIMs. **G**) The peak area ratios of PDIMs to total PIMs, lower-order PIMs, and higher-order PIMs were calculated for each *M.tb* strain and compared across drug resistance categories. Graphs are shown as M±SD, where each data point represents an individual lipid extraction. N = 12-20 extracts per drug resistance category, where the “n” value represents the number of biological replicas performed (*i.e.* the number of times a *M.tb* strain was grown and lipid extraction was conducted).. Graphs were analyzed using One-way ANOVA with post-hoc Dunnett’s test. In some cases, data sets that did not meet the assumption of normality were log transformed. Transformed data sets that still did not meet the assumption of normality were analyzed using Kruskal-Wallis nonparametric test and Dunn’s test for multiple comparisons.; *p<0.05; **p<0.01; ***p<0.001; ****p<0.0001.

We finally assessed the PDIMs:PIMs ratio to gain a greater understanding of how cell envelope hydrophobicity changes with drug resistance. TLC analysis indicated a significant increase in the PDIMs:PIMs ratio on the cell envelope of MDR- and pre-XDR-*M.tb* strains, with pre-XDR strains having the highest PDIMs:PIMs ratio when compared to DS-*M.tb* strains (**Fig. 4C**; **Suppl. Fig. S1C**). The PDIM ratio related to lower-order PIMs and higher-order PIMs were both significantly increased compared to DS-*M.tb* strains (**Fig. 4C**). Our LC-MS analysis also confirmed these TLC results, showing trends for MDR strains and significant increases in PDIMs:PIMs, PDIMs:Lower-order PIMs, and PDIMs:Higher- order PIMs ratios in pre-XDR strains compared to DS-*M.tb* strains (**Fig. 4G**; **Suppl. Fig. S1F**). Overall, these findings indicate that the DR-*M.tb* strains studied have increased levels of the hydrophobic PDIMs and decreased levels of hydrophilic higher-order PIMs on their cell envelope, suggesting that DR-*M.tb* strains might have an overall increase in cell envelope hydrophobicity compared to DS*-M.tb* strains.

### DR-*M.tb* strains have similar amounts but more surface exposure of ManLAM compared to DS-*M.tb* strains

The lipoglycans LM and ManLAM are biosynthetically associated with PIMs and are also considered virulence factors on the *M.tb* cell envelope.^48^ Since we saw significant decreases in higher-order PIMs in DR-*M.tb* using LC-MS (**Fig. 4F**), we next focused on analyzing the content of LM and ManLAM on the studied strains by using SDS-PAGE and periodic acid-silver staining (**Fig. 5**). Interestingly, we saw that there were no significant changes in the levels of LM or ManLAM across drug resistance categories (**Fig. 5A-B**; **Suppl. Fig. S4A-B**). These data indicate that while higher-order PIMs are decreased in DR-*M.tb* strains, levels of lower-order PIMs and the associated lipoglycans LM and ManLAM overall amounts seem to be conserved across drug resistance categories, even with obvious strain-specific variation within each drug resistance category.

**Figure 5.**
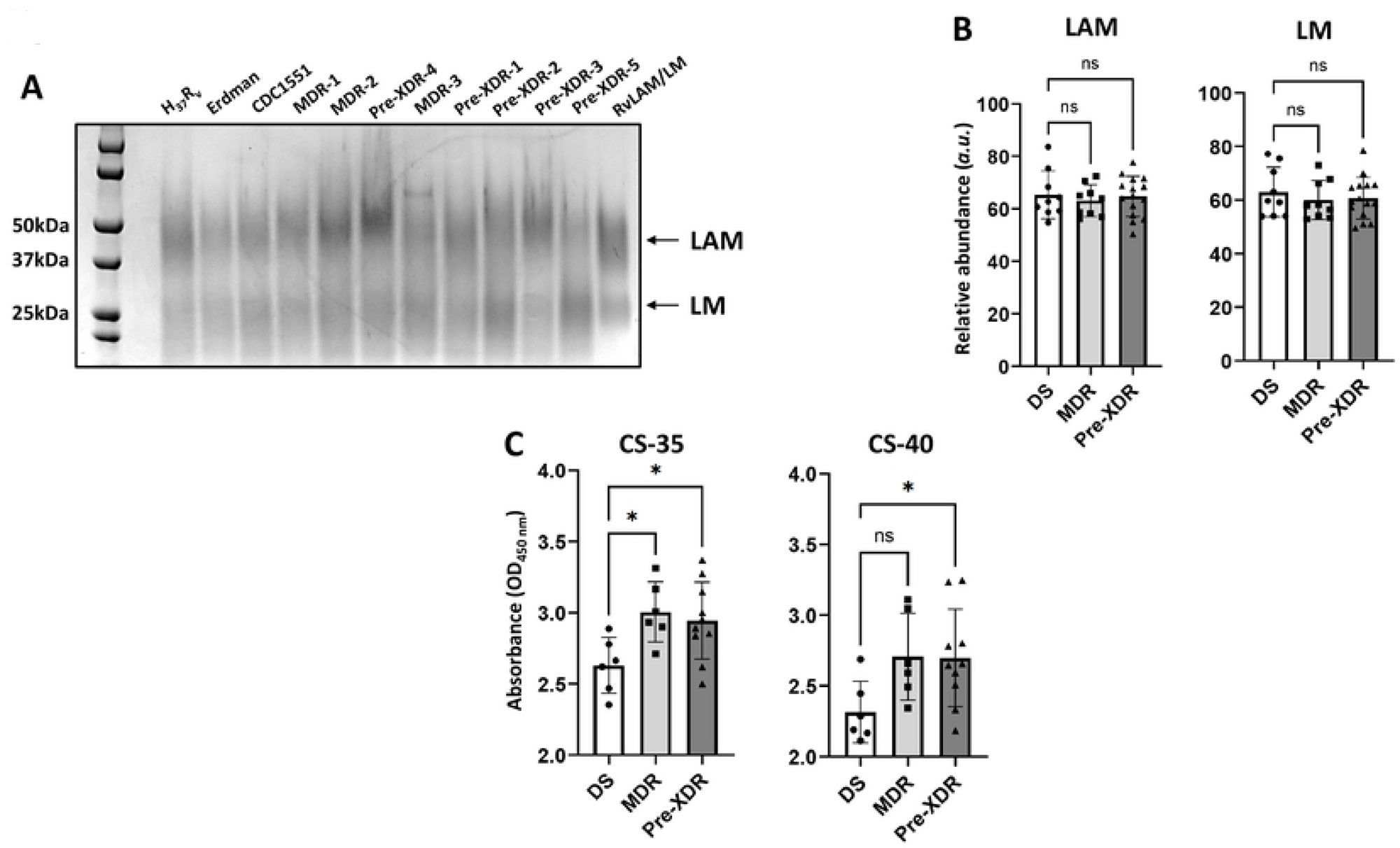
DR-*M.tb* strains have similar amounts but more surface exposure of mannose-capped lipoarabinomannan (ManLAM) compared to DS-*M.tb* strains. Lipoglycans (normalized by biomass) were extracted from each *M.tb* strain via quick LAM extraction and run using SDS-PAGE gels. **A)** Representative SDS-PAGE gel of n = 3 is shown. **B)** SDS-PAGE gels were quantified via densitometry analysis and compared across drug resistance categories for ManLAM and lipomannan (LM). Graphs are shown as M±SD, where each data point represents an individual quick LAM extraction. N = 9-15 extracts per drug resistance category, where the “n” value represents the number of biological replicas performed (*i.e.* the number of times a *M.tb* strain was grown and quick LAM extraction was conducted). **C)** Whole cell ELISAs were performed assessing the surface exposure of ManLAM and compared across drug resistance categories using monoclonal antibodies CS-35 (BEI Resources; Catalog #NR-13811) and CS- 40 (BEI Resources; Catalog #NR-13812). Graphs are shown as M±SD, where each data point represents the average value of three triplicate wells per *M.tb* strain with n=2 individual experiments performed. Graphs were analyzed using One-way ANOVA with post-hoc Dunnett’s test; *p<0.05.

Although levels of ManLAM did not differ across drug resistance categories, it is possible that the surface exposure of ManLAM may vary with drug resistance, as ManLAM may be less exposed on the surface of DR-*M.tb* that have increased PDIMs:PIMs ratios. To test this, we performed whole cell ELISAs for each *M.tb* strain using two anti-LAM mAbs, CS-35 and CS-40. Unexpectedly, we found that the surface exposure of ManLAM increased as drug resistance increased, with both MDR and pre-XDR *M.tb* strains having more ManLAM surface exposure compared to DS strains (**Fig. 5C**; **Suppl. Fig. S4C-D**). Thus, it seems that there is an increase in the surface exposure of ManLAM in DR-*M.tb* strains.

### DS- and DR-*M.tb* strains have strain-specific variability in cell envelope composition compared to the lab standard *M.tb* H_37_R_v_

While there were clear differences in the cell envelope composition across drug resistance categories, we also wanted to examine strain-specific biochemical differences. Even within DS-*M.tb* strains, we saw that *M.tb* Erdman had decreased PIMs (including lower- and higher-order PIMs), ManLAM levels, and surface exposure of ManLAM, while CDC1551 had decreased PDIMs, PDIMs:PIMs ratio, and surface exposure of ManLAM compared to H_37_R_v_ (**Table 4; Suppl. Figs. S1, 3-4**). Based on our LC-MS analysis, MDR- 1/3 and Pre-XDR-4 had very similar levels of PDIMs, but less PIMs (both lower- and higher-order PIMs) than H_37_R_v_ and thus, increased PDIMs:PIMs ratios. Additionally, MDR-2/Pre-XDR-1 both had lower levels of PDIMs, total PIMs, and higher-order PIMs compared to H_37_R_v_, ultimately leading to a decreased PDIMs:PIMs ratio in both strains. In contrast, Pre-XDR-2/3/5 all had increased PDIMs, and Pre-XDR- 2/5 had decreased PIMs (both lower- and higher-order PIMs), ultimately leading to a larger increase in their PDIMs:PIMs ratios.

**Table 4.**
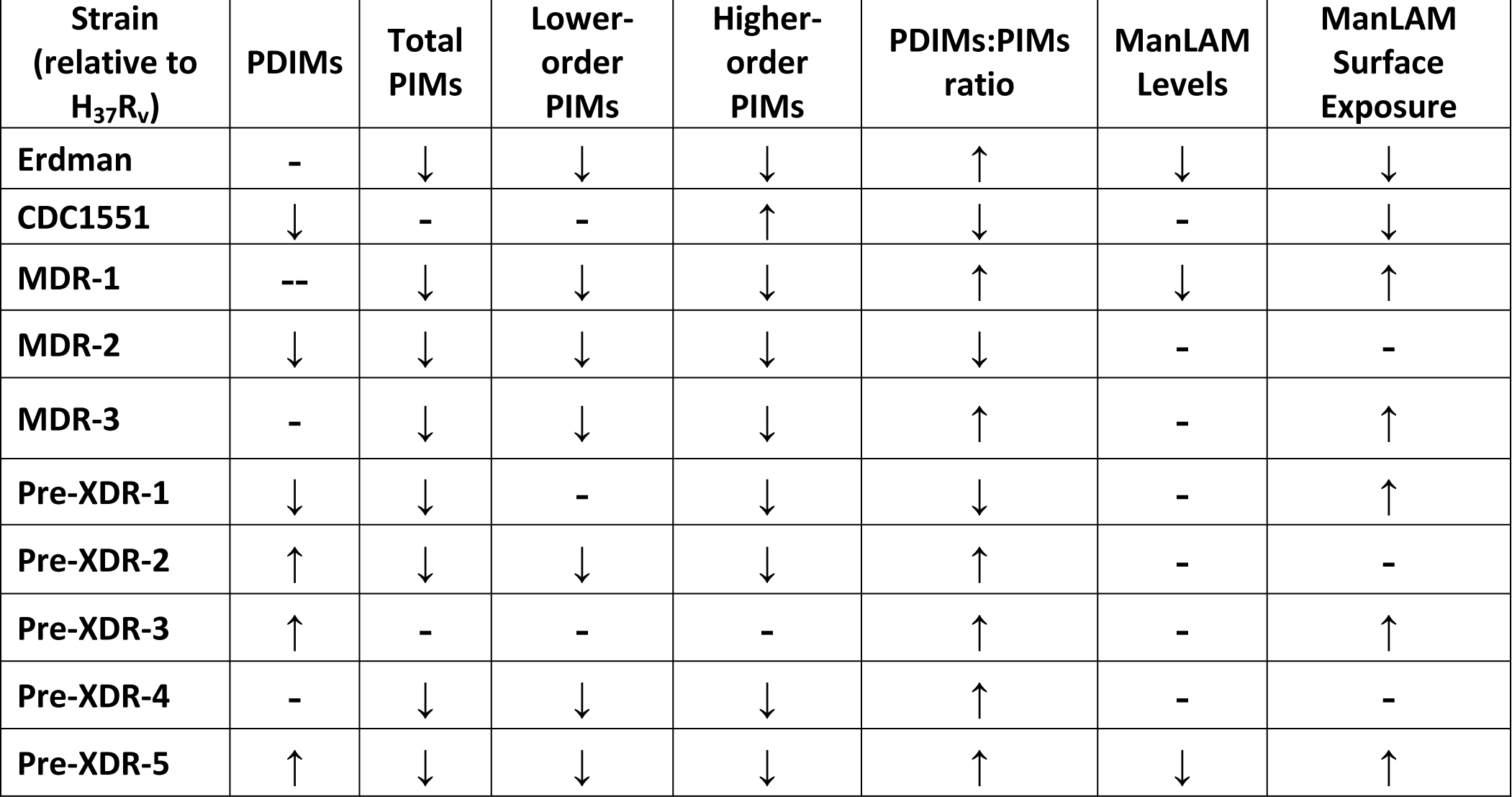
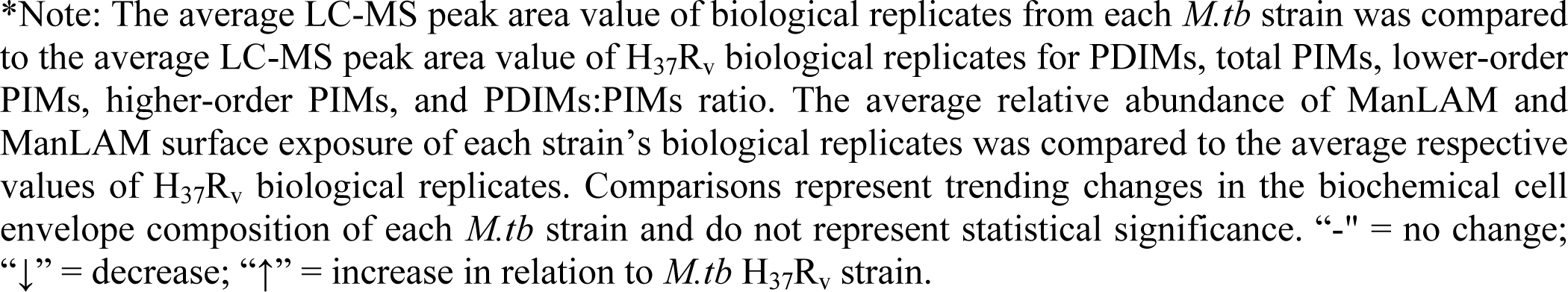
Comparison of the Biochemical Composition of Individual *M.tb* Strains*.

| Strain<br>(relative to<br>H<sub>37</sub>R<sub>v</sub>) | PDIMs | Total<br>PIMs | Lower-order<br>PIMs | Higher-order<br>PIMs | PDIMs:PIMs<br>ratio | ManLAM<br>Levels | ManLAM<br>Surface<br>Exposure |
| --- | --- | --- | --- | --- | --- | --- | --- |
| Erdman | - | ↓ | ↓ | ↓ | ↑ | ↓ | ↓ |
| CDC1551 | ↓ | - | - | ↑ | ↓ | - | ↓ |
| MDR-1 | -- | ↓ | ↓ | ↓ | ↑ | ↓ | ↑ |
| MDR-2 | ↓ | ↓ | ↓ | ↓ | ↓ | - | - |
| MDR-3 | - | ↓ | ↓ | ↓ | ↑ | - | ↑ |
| Pre-XDR-1 | ↓ | ↓ | - | ↓ | ↓ | - | ↑ |
| Pre-XDR-2 | ↑ | ↓ | ↓ | ↓ | ↑ | - | - |
| Pre-XDR-3 | ↑ | - | - | - | ↑ | - | ↑ |
| Pre-XDR-4 | - | ↓ | ↓ | ↓ | ↑ | - | - |
| Pre-XDR-5 | ↑ | ↓ | ↓ | ↓ | ↑ | ↓ | ↑ |

\*Note: The average LC-MS peak area value of biological replicates from each *M.tb* strain was compared to the average LC-MS peak area value of H<sub>37</sub>R<sub>v</sub> biological replicates for PDIMs, total PIMs, lower-order PIMs, higher-order PIMs, and PDIMs:PIMs ratio. The average relative abundance of ManLAM and ManLAM surface exposure of each strain's biological replicates was compared to the average respective values of H<sub>37</sub>R<sub>v</sub> biological replicates. Comparisons represent trending changes in the biochemical cell envelope composition of each *M.tb* strain and do not represent statistical significance. "-" = no change; "↓" = decrease; "↑" = increase in relation to *M.tb* H<sub>37</sub>R<sub>v</sub> strain.

Interestingly, we also saw strain dependent variability in ManLAM levels (**Table 4; Suppl. Fig. S4A**) and surface exposure in MDR- and pre-XDR *M.tb* strains (**Table 4; Suppl. Fig. S4C-D**). Indeed, MDR-2/3and Pre-XDR-1/2/3/4 had similar levels of ManLAM compared to H_37_R_v_, but MDR-1 and Pre-XDR-5 had slightly decreased ManLAM levels in comparison. Surface exposure of ManLAM was also varied between *M.tb* strains, with MDR-1/3 and Pre-XDR-1/3/5 all showing increased surface exposure relative to all DS-*M.tb* strains. However, MDR-2 and Pre-XDR-2/4 appeared to have similar surface exposure of ManLAM in comparison to H_37_R_v_, although they still appeared to have greater surface exposure compared to *M.tb* Erdman and CDC1551 (**Table 4**).

### DR-*M.tb* strains have decreased association with human monocyte-derived macrophages (hMDMs)

Previous studies demonstrate that *M.tb* strains with truncated (shorter) ManLAM and reduced amounts of higher-order PIMs have decreased association with primary human macrophages.^25^ Thus, it is plausible that the increased PDIMs:PIMs ratio in DR-*M.tb* strains may result in changes in *M.tb*-host interactions during infection. To test this, we infected hMDMs with each *M.tb* strain and assessed *M.tb* association (**Fig 6; Suppl. Fig. S5**). Our results indicate association of approximately 60 bacilli per 100 MDMs for DS-*M.tb* strains, but there was greater variation for DR-*M.tb*. Indeed, MDR-2/Pre-XDR-4 had approximately 30 bacilli per 100 hMDMs, Pre-XDR-1 had approximately 20 bacilli per 100 hMDMs, and MDR-1/3 and Pre-XDR-2/3/5 had approximately 10 bacilli per 100 hMDMs (**Fig 6B**). Although there was variability in association between DR-*M.tb* strains, we saw overall that there was a significant and gradual decrease in macrophage association in MDR- and pre-XDR-*M.tb* compared to DS-*M.tb* strains, indicating that association of *M.tb* with hMDMs decreases as drug resistance increases (**Fig. 6B; Suppl. Fig. S5A-B**).

**Figure 6.**
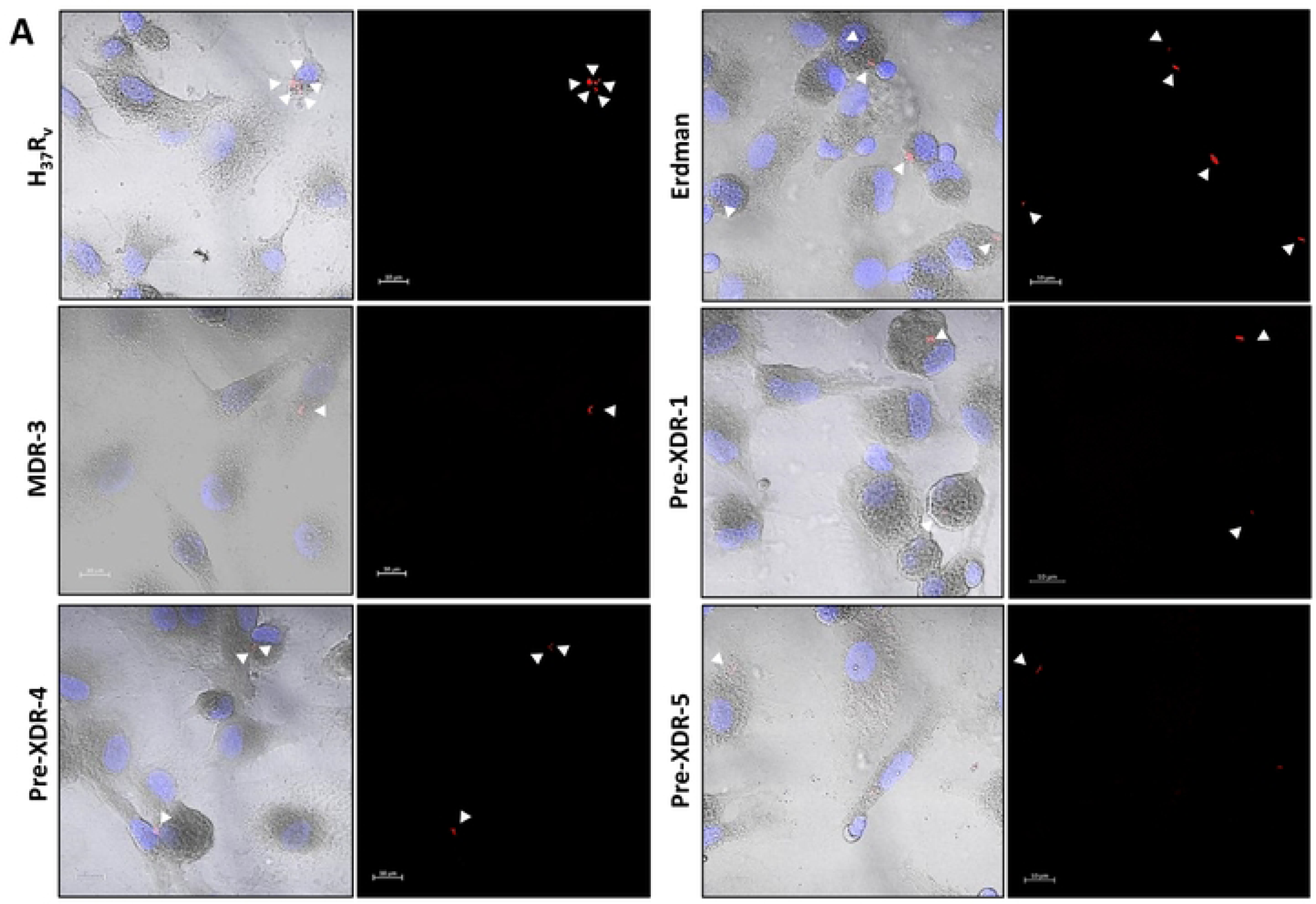
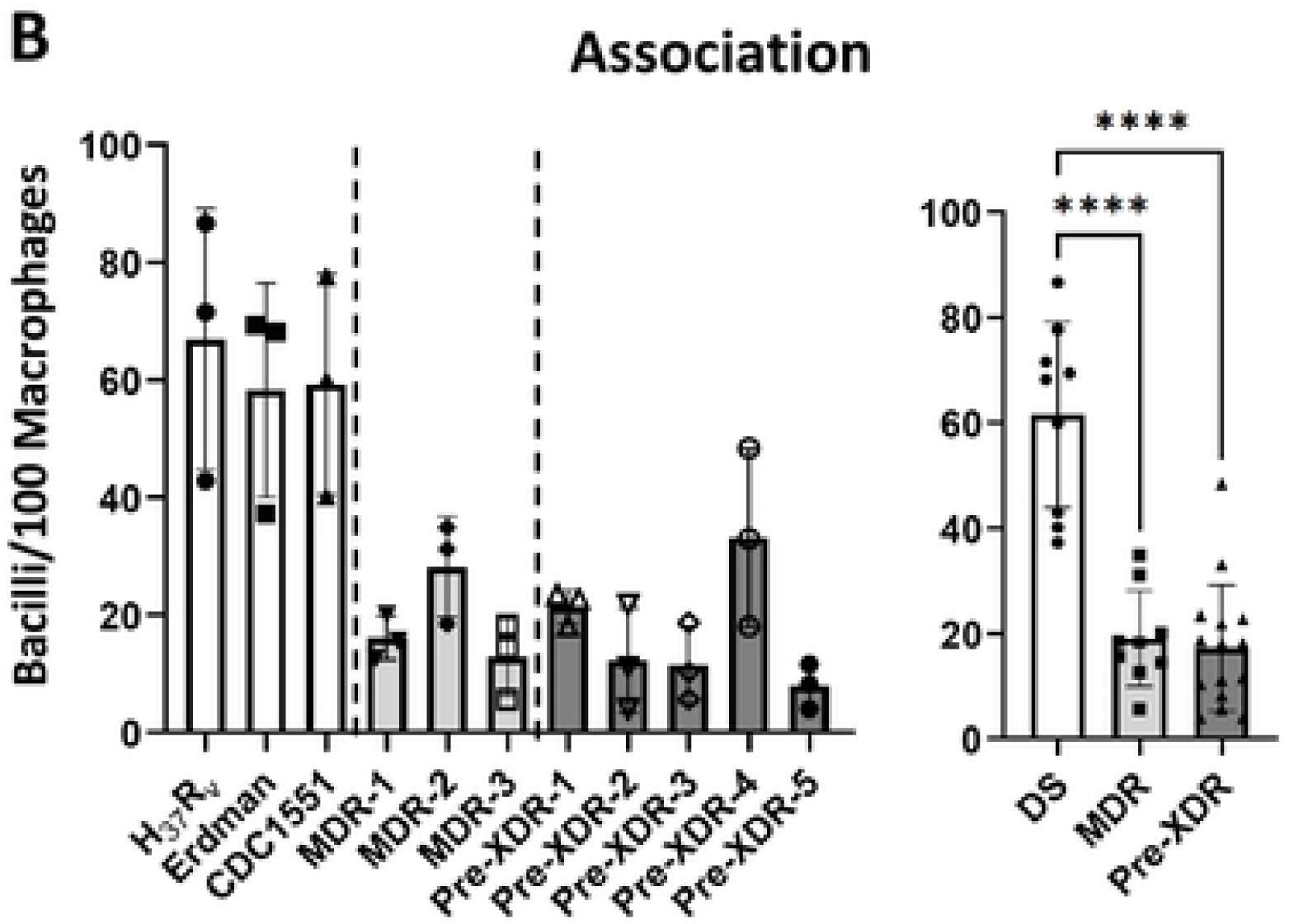
DR-*M.tb* strains have decreased association with human macrophages compared to DS- *M.tb* strains. hMDMs from human blood donors were infected with each *M.tb* strain (MOI 10:1) and assessed for association (measured by DIC and fluorescence microscopy) at 2 hpi. **A)** Representative images of MDMs infected with H_37_R_v_ or Erdman *M.tb* strains (top row), MDR-3 or Pre-XDR-4 *M.tb* strains (middle row), or Pre-XDR-4/5 *M.tb* strains (bottom row). White arrows indicate bacteria associated with a macrophage. **B)** Quantification of association was calculated as bacteria per 100 cells for each strain and across drug resistance categories. Graphs are shown as M±SD, where each data point represents an individual human blood donor (n = 3 donors) for each *M.tb* strain. Graphs were analyzed using One-way ANOVA with post-hoc Dunnett’s test; ****p<0.0001.

### The Pre-XDR-2 *M.tb* strain has decreased uptake but increased intracellular growth in human macrophages compared to *M.tb* H_37_R_v_

To dive deeper into how the PDIMs:PIMs ratio influences *M.tb* infection outcomes, we examined the uptake, bacterial burden, and intracellular growth rate of the Pre-XDR-2 *M.tb* strain *vs*. the DS reference strain H_37_R_v_ after infection of hMDMs (**Fig. 7**). We chose this pair of strains as Pre-XDR-2 *M.tb* has the highest PDIMs:PIMs ratios when compared to H_37_R_v_ by both TLC and LC-MS (**Table 4; Suppl. Figs. S1C, F** and **S6A-F**). However, Pre-XDR-2 had no difference in LM or ManLAM levels (**Table 4; Suppl. Figs. S4A-B** and **S6G-H**) or surface exposure of ManLAM (**Table 4; Suppl. Figs. S4C-D** and **S6I-J**) when compared to H_37_R_v_. Thus, we were able to investigate how an increase in the PDIMs:PIMs ratio, but not changes in ManLAM/LM levels or ManLAM surface exposure, in a DR-*M.tb* strain influences its bacterial burden and intracellular growth rate compared to the DS laboratory reference strain H_37_R_v_ strain.

**Figure 7.**
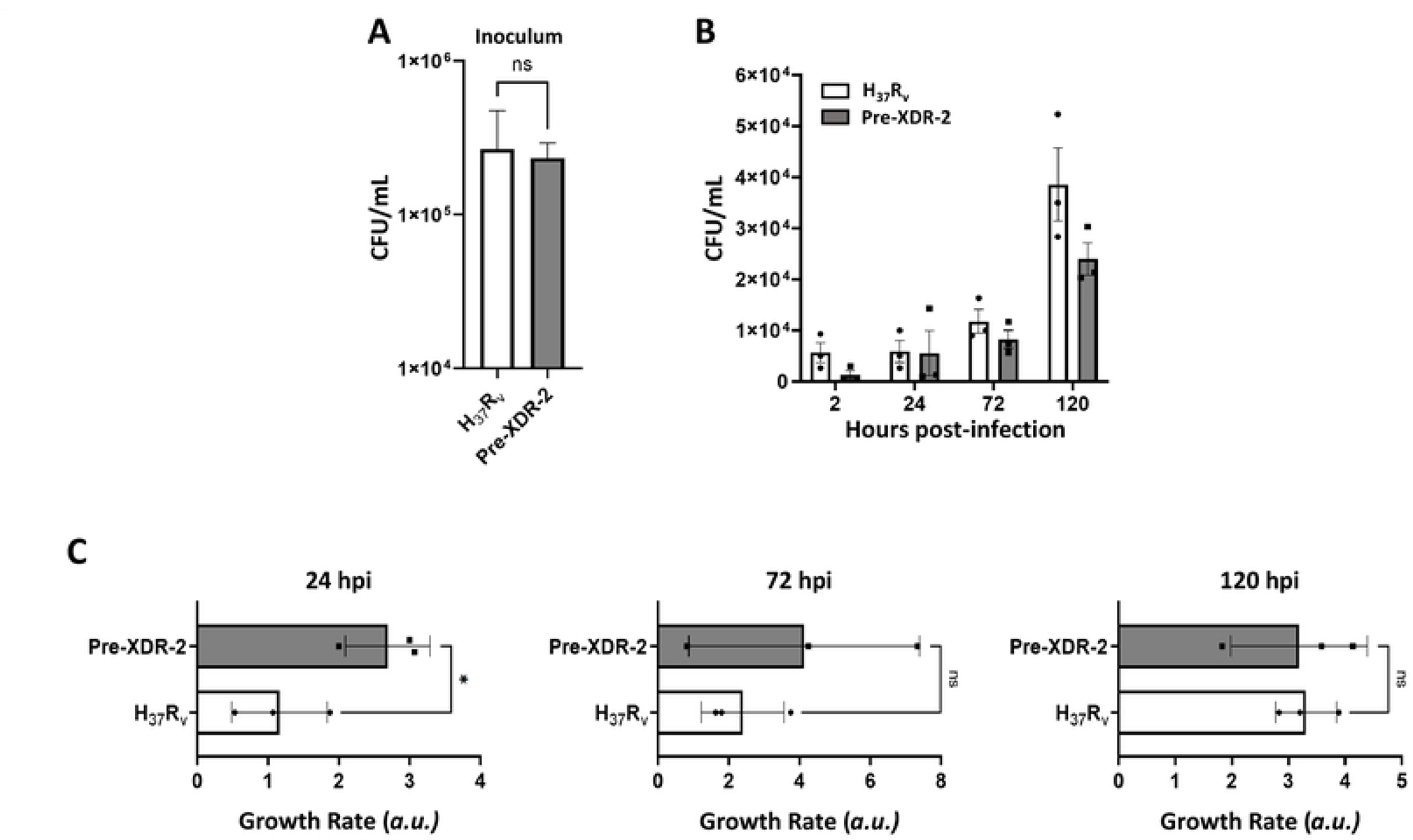
The Pre-XDR-2 *M.tb* strain has decreased uptake but an increased intracellular growth rate at early timepoints compared to H_37_R_v_. A) Representative infection inoculum showing similar amounts of bacteria in both H_37_R_v_ and Pre-XDR-2 inoculum. Representative inoculum of n=3 (performed in triplicate), M ± SD is shown. **B)** Monolayers were assessed for bacterial burden by CFU counts at 2, 24, 72, and 120 hours post-infection (hpi). Graph is shown as M±SD, where each data point represents the average value of three triplicate wells of MDMs from an individual blood donor (n = 3) for each *M.tb* strain. **C)** Growth rates for H_37_R_v_ and Pre-XDR-2 were calculated at 24, 72, and 120 hpi by taking the mean triplicate value at each timepoint and dividing it by the mean triplicate value at the previous timepoint. Graphs are shown as M±SD, where each data point represents the average value of three triplicate wells of MDMs from an individual blood donor (n = 3) for each *M.tb* strain. Graphs were analyzed using Student’s t test at each timepoint; *p<0.05.

In line with our association results, we saw that even though hMDMs were infected with similar amounts of bacteria (**Fig. 7A**), *M.tb* Pre-XDR-2 had decreased uptake at 2 hpi compared to *M.tb* H_37_R_v_ (**Fig. 7B**). Additionally, we also saw that there was a trending decrease in the bacterial burden of hMDMs infected with *M.tb* Pre-XDR-2 when compared to *M.tb* H_37_R_v_ at multiple timepoints after infection (**Fig. 7B**). Interestingly, however, when compared to *M.tb* H_37_R_v_, we observed that the intracellular growth rate of *M.tb* Pre-XDR-2 was significantly increased in hMDMs at early timepoints after infection, although this difference disappeared over time (**Fig. 7C**).

## DISCUSSION

In recent decades, DR-*M.tb* strains have emerged that pose a risk to worldwide public health. Previous studies suggest that a deficiency in the hydrophobic cell envelope lipid PDIMs results in increased cell envelope permeability and greater susceptibility of mycobacteria to anti-TB drugs.^3, 11–13^ However, there are very few studies indicating increased levels of PDIMs in DR-*M.tb* strains.^15^ Here we biochemically examined the cell envelope lipid composition of 11 *M.tb* strains that range from DS (including H_37_R_v_, Erdman, and CDC1551) to MDR and pre-XDR. We established that in this small set of *M.tb* strains, there is an increase in hydrophobic PDIMs and a decrease in hydrophilic PIMs (in particular, higher-order PIMs) on the cell envelope of the MDR- and pre-XDR-*M.tb* strains. We also presented data suggesting that levels of PIMs-associated lipoglycans LM and ManLAM are conserved across drug resistance categories, although there is increased surface exposure of ManLAM as the drug resistance of our strains increased. Finally, we provided data to support that the DR-*M.tb* strains examined have decreased association with primary hMDMs compared to DS-*M.tb* strains, and that a DR-*M.tb* strain with increased PDIMs:PIMs ratio (*e.g.*, Pre-XDR-2) but similar ManLAM/LM levels and ManLAM surface exposure has decreased uptake, balanced by an increased intracellular growth in hMDMs early post-infection which stabilized over time in relation to the laboratory standard *M.tb* H_37_R_v_ strain. Importantly, from the *M.tb* strains studied, the DR-*M.tb* strains seem to have PDIMs isomers that were not observed for the DS-*M.tb* strains. These observations support the notion that DR-*M.tb* strains may have an altered, more hydrophobic cell envelope composition that could ultimately influences *M.tb*-host interactions during infection (**Fig. 8**).

**Figure 8.**
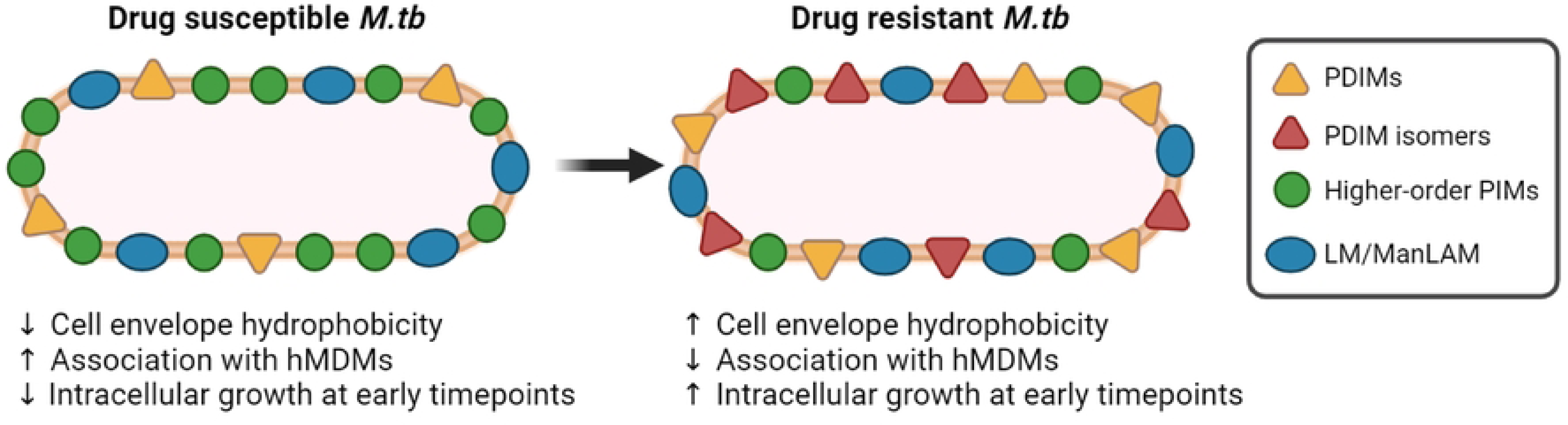
Schematic representation of the main findings in this study. Based on our results, as drug resistance increases, there seems to be an overall increase in the hydrophobicity of the *M.tb* cell envelope. This increase in hydrophobicity is most noticeable by the increase in hydrophobic PDIMs (including PDIM isomers not present in DS strains) and decrease in hydrophilic higher-order PIMs in DR-*M.tb* strains. Moreover, DR-*M.tb* strains also have decreased association with hMDMs but an increased intracellular growth rate at early timepoints after hMDM infection. Overall, our findings point towards an increase in the hydrophobicity of the DR-*M.tb* cell envelope that may influence early *M.tb*-host cell interactions during infection. This illustration was created with BioRender (https://biorender.com/). PDIMs = phthiocerol dimycocerosates; PIMs = phosphatidyl-*myo*-inositol mannosides; LM = lipomannan; ManLAM = mannose-capped lipoarabinomannan; hMDMs = human monocyte-derived macrophages; *M.tb* = *Mycobacterium tuberculosis*; “↑” indicates an increase; “↓” indicates a decrease.

This study provided many interesting observations related to variability of the *M.tb* cell envelope in the context of drug resistance. However, there are clear limitations of this work that require future studies. First, due to our use of *M.tb* clinical isolates as opposed to isogenic strains, we cannot assume that the observed changes are causally associated with an increase in drug resistance. We also cannot know at this point if one specific mutation conferring resistance to an anti-TB drug is related to the increase in PDIMs observed, or if it is the sum of several mutations defined within a drug resistance category that drives the PDIMs phenotype observed in the strains studied. Although our current observations suggest a potential link between drug resistance and increased PDIMs levels, we must confirm these observations by biochemically examining isogenic or clonal strains that differ in resistance (*e.g.*, evaluation of a lab strain or a clinical isolate that goes from pan-susceptible, to mono-resistant, to MDR, etc.). Moreover, we only focused on DR-*M.tb* clinical isolates from lineages 2 and 4, and did not observe differences in the PDIMs:PIMs ratio between lineages. However, we only examined DS-*M.tb* strains from lineage 4 and thus, it will be important to expand our set of *M.tb* strains to encompass multiple lineages in both pan- susceptible and DR strains to determine if the observed differences are associated with different lineages. Ongoing experiments in our laboratories are focused on addressing these limitations.

Nonetheless, mycobacterial PDIMs are found only in pathogenic species and are hydrophobic molecules associated with cell envelope impermeability. An initial study showed that PDIMs reduce *M.tb* cell envelope permeability of hydrophobic probes and detergents,^10^ which was confirmed in a later report.^49^ Subsequent studies went on to examine the relationship between PDIMs and drug resistance, ultimately showing that PDIMs deficiencies are associated with increased susceptibility of mycobacterial species (including *M. marinum*, *M. bovis* BCG, and *M.tb*) to multiple anti-TB drugs including RIF.^11–13^ In addition, RIF-acquired resistance has been linked to increased PDIMs levels^15^ and upregulation of genes related to PDIMs biosynthesis.^14^ These data combined suggest that PDIMs may be associated with resistance of *M.tb* to anti-TB drugs; however, there is currently a gap in knowledge related to how PDIMs levels change with drug resistance. In this study, using 11 different *M.tb* strains we present observations that DR-*M.tb* strains display a range of PDIMs levels, with pre-XDR-*M.tb* strains examined producing the most PDIMs compared to DS-*M.tb*. Our data examine the relationship of PDIMs and drug resistance from the other side of the coin, and supports previous hypotheses that PDIMs may play a role in *M.tb* cell envelope impermeability to anti-TB drugs. Our results are also in line with a previous study showing increased PDIMs levels in strains with RIF-resistance,^8^ however we found that the pre-XDR *M.tb* strains studied had increased PDIMs even compared to the MDR-*M.tb* strains that also harbored RIF resistance. Our biochemical results, in addition to the fact that our MDR- and pre-XDR-*M.tb* strains have similar genotypic drug resistance profiles within each drug resistance category, indicate that there may not be one specific anti-TB drug associated with the increased PDIMs:PIMs ratio observed; however, we did see the starkest increase in PDIMs:PIMs ratio in the majority of pre-XDR-*M.tb* strains tested (except for Pre- XDR-1). Thus, moving forward it may be beneficial to ascertain whether resistance to moxifloxacin (MFX) and/or amikacin (AMK) is associated with an increased PDIMs:PIMs ratio. It is possible that while RIF resistance may be somewhat related to increased PDIMs, additional resistance to other anti-TB drugs such as MFX or AMK may also be associated with higher levels of PDIMs. On the other hand, we may find in future studies that it is a combination of *M.tb*-host interactions during the initial infection, genetic changes, and/or exposure to multiple anti-TB drugs unique to each *M.tb* strain that influences whether there is an overall increase in cell envelope hydrophobicity.

Our study also uncovered isomers of multiple PDIM alkylforms present only in DR-*M.tb* strains. Our MS/MS spectra displayed a similar, but not identical, fragmentation pattern for the *M.tb* Pre-XDR-2 PDIM that eluted at 27.053 min. Fragmentation pattern analysis suggests that the structure of this isomer has a 29:0/29:0 mycocerosic acid composition. Additionally, substitution of a hydroxyl group for the methoxy group located on a C35 phthiocerol core would explain the same MS spectra, but with slight variations in the MS/MS fragmentation patterns. Future studies involving advanced analytical techniques such as ion mobility MS will allow us to resolve these isomers and determine their exact structures.

While there were strain-specific variations in detection and abundance of these PDIM isomers, it appeared that they were more present in PDIM A alkylforms compared to PDIM B for the majority of our *M.tb* strains (except for MDR-2 and Pre-XDR-5). Indeed, additional isomers of PDIM A alkylforms appeared to make up most of its peak area, while additional isomers of PDIM B alkylforms only made up a fraction of the peak area (or were not detected at all) for the majority of the clinical isolates studied. MS fragmentation of the isomers demonstrates that those with a later LC retention time represent PDIM species based on phthiocerol while those with an earlier retention time were based on the less commonly observed phthiotriol. The final stage of phthocerol biosynthesis converts the phthiodiolone to phthiotriol via the ketoreductase Rv2951c, and then the phthiotriol to phthiocerol via the methyltransferase Rv2952.^50^ Huet *et al.* previously demonstrated the accumulation of phthiotriol based PDIM structures in the lineage 2 “*Beijing*” strains of *M.tb.* This was attributed to a point mutation in *rv2952* that reduced its methyltransferase activity.^50^ Importantly, the DR-*M.tb* strains that produced the phthiotriol based PDIM isomers in this study included lineage 2 and 4 strains, suggesting that DR lineage 4 strains may also have reduced activity of the Rv2952 methyltransferase. Future studies are required to determine whether these strains possess a similar mutation in *rv2952* or if another mechanism is responsible for this phenotype. Moreover, because our study was limited to only 11 *M.tb* strains, future work examining a larger subset of DS- and DR-*M.tb* strains will be necessary to determine whether these isomers directly impact DR phenotypes.

Unlike PDIMs, hydrophilic PIMs and associated lipoglycans LM and ManLAM have been more widely studied in past years and are considered important virulence factors on the *M.tb* cell envelope. These mannose-containing glycolipids and lipoglycans can vary by strain,^23–26^ with truncated forms of ManLAM being associated with decreased phagocytosis and lower association with host cell receptors such as the MR.^25, 26^ Additionally, our previous work suggests that decreased amounts of higher-order PIMs (also recognized by the MR^51^) are also related to decreased association with primary human macrophages.^25^ One previous study examined an ethambutol-resistant *M.tb* strain and found that it made a largely heterogeneous population of LAM;^26^ however, there is limited data on how levels of PIMs and their associated lipoglycan change in relation to drug resistance. In our study, we provide data suggesting that while amounts of lower-order PIMs, LM, and ManLAM are similar across the strains in each drug resistance category, there is a significant decrease in higher-order PIMs in our MDR- and pre-XDR-*M.tb* strains compared to DS-*M.tb* strains. These data indicate that there is only partial conservation of the PIMs biosynthetic pathway in the DR-*M.tb* strains we examined, with the focus being on production of LM and ManLAM rather than higher-order PIMs. In addition, our association results are in line with those of previous studies showing that decreases in higher-order PIMs result in decreased association of *M.tb* with primary human macrophages. These data, combined with our biochemical results showing decreased higher-order PIMs in all DR-*M.tb* strains (except for Pre-XDR-3) suggest that a decrease in higher-order PIMs specifically may be associated with decreased association of *M.tb* with MDMs, and that as higher- order PIMs decrease as drug resistance increases, so does association with human macrophages. Our previously published work demonstrates that *M.tb* strains with less exposed ManLAM and reduced higher-order PIMs but abundant PDIMs, triglycerides, and phenolic glycolipids have decreased phagocytosis but faster intracellular growth in human macrophages compared to strains with more complex ManLAM and increased amounts of higher-order PIMs such as H_37_R_v_ and Erdman.^25^ While this work underscores the importance of the cell envelope composition in *M.tb*-host interactions, it does not address the topic of drug resistance. We observed a similar phenomenon in our current work in the context of *M.tb* drug resistance. We saw that the MDR- and pre-XDR-*M.tb* strains we examined have decreased higher-order PIMs along with decreased association with human macrophages. By assessing intracellular growth via CFUs, we also saw that the Pre-XDR-2 *M.tb* strain with the highest PDIMs:PIMs ratio (but similar levels and surface exposure of ManLAM) has decreased uptake at 2 hpi compared to the H_37_R_v_ strain. Additionally, our data show a similar pattern where Pre-XDR-2 *M.tb* had an increased intracellular growth rate compared to *M.tb* H_37_R_v_ at early timepoints post-infection; however, this differential growth rate waned over time. Future studies will be required to examine how *M.tb* strains in each drug resistance category interact with host cells, and how changes in these interactions influence the immune response during infection.

Altogether, our biochemical results provide a compelling observation suggesting that DR-*M.tb* strains may have increased levels of hydrophobic PDIMs and decreased levels of hydrophilic PIMs defining a differential PDIMs:PIMs ratio that increases as the drug resistant profile increases (*e.g.* from DS to MDR to pre-XDR), and that changes in these cell envelope virulence factors result in decreased association but an enhanced intracellular growth rate in hMDMs at early timepoints post-infection (**Fig. 8**). Our study highlights a potential link between *M.tb* cell envelope lipids and drug resistance, and how changes in these lipids may ultimately influence DR-*M.tb* infection outcomes. Future studies expanding the categories of *M.tb* strains to include RIF-resistant (RR) and XDR categories, as well as different genotypic lineages and increased number of strains in each drug resistance category, will be crucial to map exactly how the *M.tb* cell envelope changes as drug resistance increases. This knowledge will ultimately further our understanding of DR-TB and pave the way to developing novel anti-TB drugs and diagnosis tools to eradicate this deadly disease.

## ACKNOWLEDGEMENTS

We acknowledge the Texas Biomedical Research Institute Biosafety Level 3 (BSL-3) Operations Program for their services and support.

## SUPPLEMENTAL MATERIAL

The online version of this article contains supplemental material.

## AUTHOR CONTRIBUTIONS

Study design and conceptualization, A.S. and J.B.T.; Provision of materials, B.K. and B.M.; Experimental procedures, A.S., M.N.I., M.W., and A.H.; Acquisition of data and formal analyses, A.S. and M.N.I.; Writing—original draft, A.S. and J.B.T.; Visualization and BioRender figures, A.S.; Writing—review and editing, B.K., B.M., J.T.B., and J.B.T.; Funding acquisition, A.S. and J.B.T. All authors have read and agreed to the published version of the manuscript.

